# Serine arginine-rich splicing factor (SRSF7) cooperates with the histone methyltransferase KMT5a to promote the type I interferon response via transcriptional activation of IRF7

**DOI:** 10.1101/2023.05.09.540055

**Authors:** Haley M. Scott, Mackenzie H. Smith, Aja K. Coleman, Summer L. Apostalo, Allison R. Wagner, Robert O. Watson, Kristin L. Patrick

## Abstract

Tight regulation of macrophage immune gene expression is required to fight infection without risking harmful inflammation. The contribution of RNA binding proteins (RBPs) to shaping the macrophage response to pathogens remains poorly understood. Transcriptomic analysis revealed that a member of the serine/arginine-rich (SR) family of mRNA processing factors, SRSF7, is required for optimal expression of a cohort of interferon stimulated genes (ISGs) in macrophages. Using genetic and biochemical assays, we discovered that in addition to its canonical role in regulating alternative splicing, SRSF7 drives transcription of interferon regulatory transcription factor 7 (IRF7) to promote antiviral immunity. At the *Irf7* promoter, SRSF7 maximizes STAT1 transcription factor binding and RNA polymerase II elongation via cooperation with the H4K20me1 histone methyltransferase KMT5a (SET8). These studies define an unorthodox role for an SR protein in activating transcription and reveal an unappreciated RNA binding protein-chromatin network that orchestrates macrophage antiviral gene expression.

## INTRODUCTION

Macrophages (MΦ) are innate immune cells that serve as the body’s first line of defense against pathogens. They express a panoply of pattern recognition receptors that detect pathogen- and damage-associated molecular patterns (PAMPs and DAMPs) and transmit these “danger signals” to the nucleus though complex signal transduction cascades. Generally, these pathogen sensing cascades terminate with phosphorylation and nuclear translocation of specialized immune transcription factors (e.g. NFκB, IRF3, IRF7). Because innate immune transcription factors exert control over hundreds of genes with massive inflammatory potential, their misregulation can have dire consequences for a host: innate responses that are too weak enable pathogen survival and replication, whereas responses that are too strong can trigger a cytokine storm, chronic inflammation, and autoimmunity. While the mechanisms through which innate immune transcription factors are regulated post-translationally—via phosphorylation, dimerization, and nuclear translocation—are well-established (Caamano and Hunter, 2002; Jefferies, 2019; Ning et al., 2011; Smale, 2012), how cells control expression of these master regulators at the transcriptional and post-transcriptional levels remains less well-understood.

A growing literature has built a strong case for RNA binding proteins (RBPs) in shaping the transcriptomes of specialized cells like macrophages. SR (serine-arginine rich) proteins are a 15-member family of RBPs that are broadly defined as regulators of alternative splicing. By virtue of distinct RNA recognition motifs, SRs bind to unique exonic splicing enhancers to help direct the spliceosome to cis-splicing signals in target transcripts (Wagner and Frye, 2021). While best known for their role in pre-mRNA splicing, SRs can participate in virtually any step of RNA processing, from transcription all the way to translation (Jeong, 2017). These studies of SR function have mostly been carried out in genetically tractable “model” cell lines (e.g. K562, HEK293T, and HepG2), but we are beginning to appreciate specialized roles for SR proteins in immune cells. SRSF1 is required for regulatory T cell homeostasis and function (Katsuyama and Moulton, 2021) and several SR proteins have been implicated in T cell development via controlling alternative splicing of CD45 (Lemaire et al., 1999; ten Dam et al., 2000; Wang et al., 2001). We recently implicated SRSF6 in maintaining mitochondrial homeostasis and antiviral immunity in macrophages by controlling alternative splicing of the pro-apoptotic factor BAX (Wagner et al., 2022).

Intriguingly, differential phosphorylation of SR proteins has been reported in macrophages infected with intracellular pathogens like the bacterium *Mycobacterium tuberculosis* (Budzik et al., 2020) and the fungal pathogen *Cryptococcus neoformans* (Pandey et al., 2017). As phosphorylation of SR proteins regulates their localization and function (Misteli et al., 1998; Shin et al., 2004; Zhong et al., 2009; Zhou and Fu, 2013), these macrophage phosphoproteomics studies suggest that functionalization of SR proteins can occur downstream of pattern recognition receptor engagement. We hypothesized that these SR proteins play privileged roles in tuning the macrophage response to pathogens. Indeed, we found that stable knockdown of several SR proteins in RAW 264.7 macrophage-like cells (RAW MΦs) profoundly altered the transcriptome in resting and activated macrophages (4h post-*Salmonella* Typhimurium infection) (Wagner et al., 2021). The absence of one of these proteins, SRSF7, generated some particularly interesting phenotypes. SRSF7 (formerly known as 9G8) is a canonical member of the SR protein family. It encodes an RNA recognition motif (RRM) and an eponymic SR domain. It is the only SR family protein to also encode a CCHC-type Zn knuckle domain, which is thought to contribute to its RNA-binding specificity (Cavaloc et al., 1999; Cavaloc et al., 1994; Wang et al., 2021)). To date, its function is best studied in the context of controlling its own expression levels via poison cassette exon inclusion (Konigs et al., 2020), in regulating mRNA export (Muller-McNicoll et al., 2016), and in alternative splicing (Kadota et al., 2020). Importantly, altered expression of SRSF7 is associated with a variety of human cancers (Boguslawska et al., 2016; Cun et al., 2021; Fu and Wang, 2018; Song et al., 2020; Xu et al., 2022).

In RAW MΦs lacking SRSF7, we identified a large cohort of genes whose expression relied on SRSF7 at rest and after *Salmonella* infection. Curiously, in both conditions, many SRSF7-impacted genes fell into the category of interferon stimulated genes (ISGs). After computational quantification of alternatively spliced genes ruled out a role for SRSF7 in mediating alternative splicing of ISG pre-mRNAs, we set out to elucidate the mechanism though which SRSF7 controls the type I IFN response in macrophages. Herein, we report a novel role for SRSF7 in controlling the transcription of interferon responsive factor 7 (IRF7), a critical player in activating the type I IFN response (Ning et al., 2011). Specifically, we demonstrate that SRSF7 enables recruitment of signal transducer and activator of transcription 1 (STAT1) to the *Irf7* promoter and relieves RNA polymerase II pausing. SRSF7 accomplishes this through interactions with the histone methyltransferase KMT5a (a.k.a. SET8), which deposits H4K20me1 at the *Irf7* promoter following LPS treatment to maximize STAT1 association. By revealing a complex regulatory network at the *Irf7* promoter that controls homeostatic and pathogen-induced type I interferon signaling, our work identifies an unexpected mechanism through which SR proteins can orchestrate macrophage innate immune gene expression and dictate infection outcomes.

## RESULTS

### SRSF7-dependent alternative splicing is independent of macrophage activation state

As differential phosphorylation of SRSF7 was detected over the course of *M. tuberculosis* infection in macrophages (Budzik et al., 2020), we were motivated to examine how macrophage activation might impact its function. To test the hypothesis that alternative splicing (AS) events might be distinct between resting and activated macrophages, signifying functionalization of SRSF7, we generated two *Srsf7* knockdown cell lines by transducing RAW MΦs (*Srsf7* KD #1 and *Srsf7* KD #2) with lentivirus containing shRNA hairpin constructs designed against the *Srsf7* 3’UTR and selected these cell lines alongside a scramble (SCR) control (**Fig. 1A-B**). RNA-seq was performed comparing SCR and *Srsf7* knockdown (KD) #2 in two conditions: uninfected (UN) or infected with the gram-negative bacterial pathogen *Salmonella enterica* serovar Typhimurium (4h post-infection; +SAL). *Salmonella* induces robust innate immune responses in macrophages though TLR4, which activates NFκB downstream of the MyD88 adapter and IRF3/IRF7 downstream of TRIF (Fitzgerald et al., 2003). Differential expression analysis was conducted using the Qiagen CLC Genomics Workbench and quantification of alternative splicing changes was carried out using MAJIQ (Modeling Alternative Junction Inclusion Quantification) (Vaquero-Garcia et al., 2016). Consistent with SRSF7’s designation as a splicing factor and previous studies implicating it in alternative splicing specific pre-mRNAs (Gu et al., 2012; Kadota et al., 2020), we identified 1857 total SRSF7-dependent alternative splicing events in resting RAW MΦs and 1909 SRSF7-dependent events in *Salmonella*-infected RAW MΦs (4h post-infection) (**Fig. 1C**) (p<0.05; >10% PSI) (**Supplemental Table 1**). As expected for an SR protein that promotes exon recognition/inclusion, most changes were at the level of exon skipping. Of the 964 genes subject to SRSF7-dependent alternative splicing in resting MΦs and the 993 genes in *Salmonella*-infected macrophages, 493 of these genes overlapped (**Fig. 1D**). Notably, only a handful of the detected alternative splicing changes occurred in genes that were also differentially expressed in *Srsf7* knockdown macrophages, suggesting that *Srsf7* impacts steady state RNA levels in a splicing-independent fashion (**Fig. 1E**).

**Figure 1:**
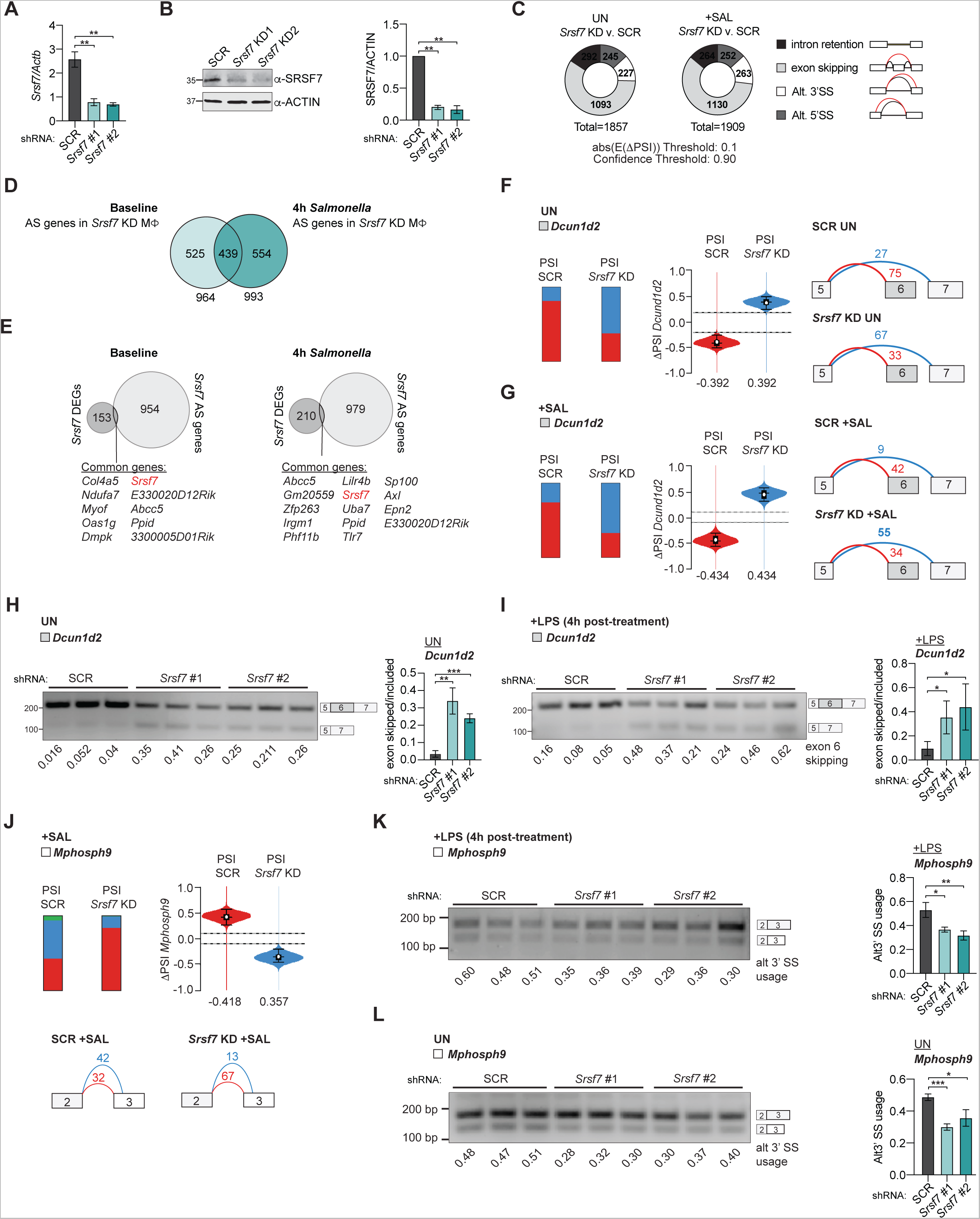
Macrophage activation does not regulate SRSF7 control of alternative splicing. **A.** RT-qPCR of *Srsf7* in *Srsf7* KD RAW MΦs relative to SCR controls. **B.** Immunoblot of SRSF7 in shSCR, sh*Srsf*7 #1, sh*Srsf*7 #2 RAW MΦs. Right, quantification. **C.** Local splicing variations (AS) in *Srsf7* RAW MΦs compared to SCR at baseline (UN; left) and +*Salmonella* Typhimurium (4h) (+SAL; right). ΔPSI > 0.1 confidence threshold set at 0.90. **D.** Venn diagram of genes containing one or more AS event in UN vs. +SAL samples. **E.** Venn diagram of differentially expressed genes (DEGs) vs. alternatively spliced genes (AS) in UN and +SAL samples. **F.** MAJIQ PSI quantification of junctions with SCR (left) and *Srsf7* KD (right) in UN samples. Violin plot of ΔPSI *Dcun1d2* SRSF7-dependent local splicing variation. Splice graph of *Dcun1d2* in SCR (top) and *Srsf7* KD (bottom) generated by MAJIQ/VOILA **G.** As in F but for +SAL samples. **H.** Semi-quantitative RT-PCR of *Dcun1d2* in SCR, *Srsf7* KD #1 and KD #2 RAW MΦs. Each sample in biological triplicate. Right, quantification. **I.** As in H but for *Dcun1d2* +LPS (4h) **J.** As in F but *Mphosph9.* **K.** As in H but *Mphosph9* (+4h LPS). **L.** As in H but *Mphosph9* in UN samples. Data are expressed as a mean of three or more biological replicates with error bars depicting SEM. Statistical significance was determined using two tailed unpaired student’s t test. *=p<0.05, **=p<0.01, ***=p<0.001.

**Figure S1:**
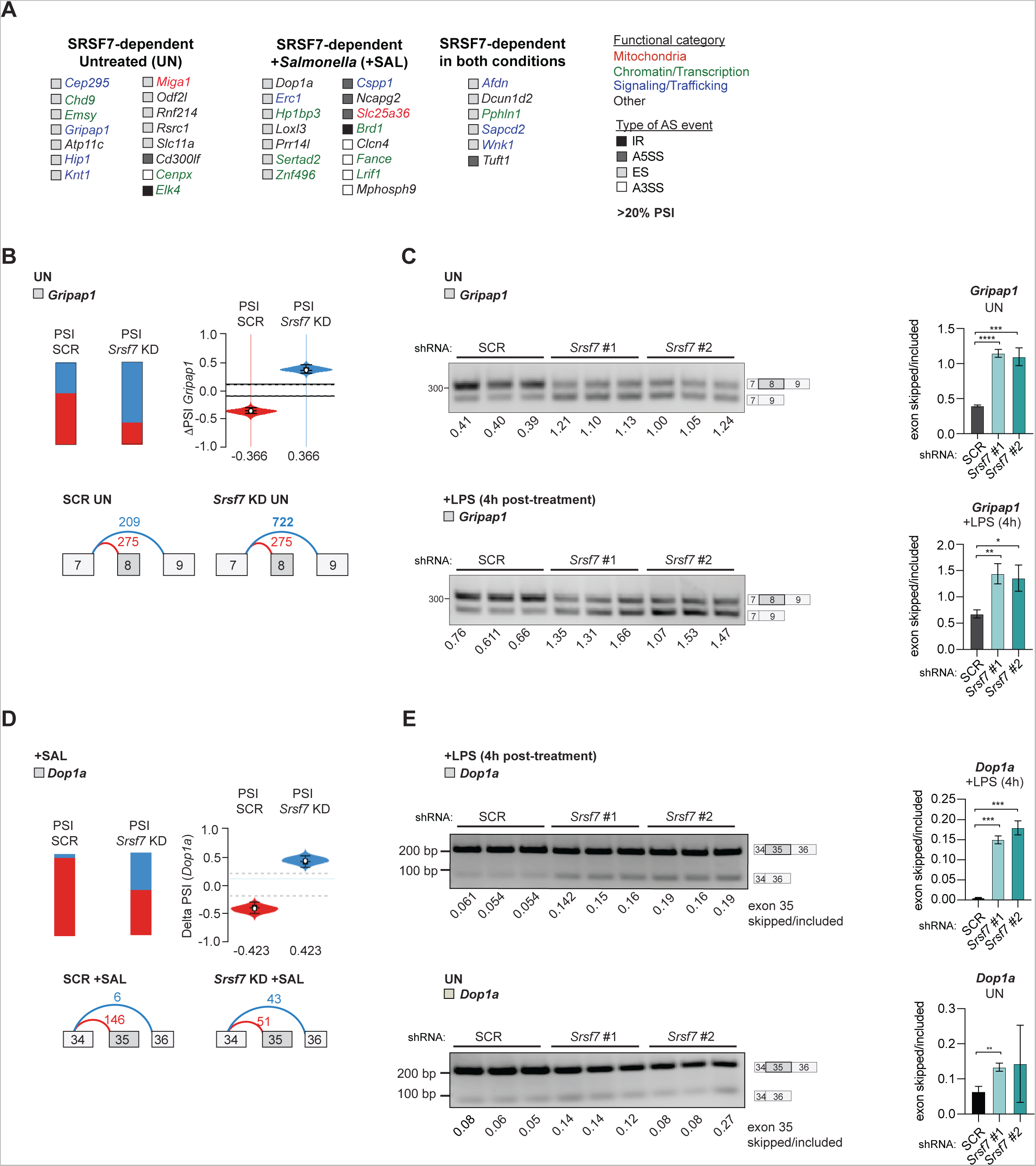
Supplemental data for Figure 1. **A.** Functional categorization of alternatively spliced transcripts in *Srsf7* knockdown RAW MΦs vs. SCR (UN, +SAL, and ΔPSI > 0.2 confidence threshold set at 0.95. **B.** MAJIQ PSI quantification of *Gripap1* junctions with SCR (left) and *Srsf7* KD (right). Violin plot of ΔPSI *Gripap1* SRSF7-dependent local splicing variation. Splice graph of *Gripap1* in SCR (bottom left) and *Srsf7* KD (bottom right) generated by MAJIQ/VOILA at baseline. **C.** Semiquantitative PCR of *Gripap1* in SCR, *Srsf7* KD1 and KD2 RAW MΦs, UN (top) and 4h +LPS (bottom). Each sample in biological triplicate. Right, quantification. **D.** MAJIQ PSI quantification of *Dop1a* junctions with SCR (left) and *Srsf7* KD (right). Violin plot of ΔPSI *Dop1a* SRSF7-dependent local splicing variation. Splice graph of *Dop1a* in SCR (bottom left) and *Srsf7* KD (bottom right) generated by MAJIQ/VOILA at baseline. **E** Semiquantitative PCR of *Gripap1* in SCR, *Srsf7* KD #1 and KD #2 RAW MΦs, 4h +LPS (top) and UN (bottom). Each sample in biological triplicate. Right, quantification. Data are expressed as a mean of three biological replicates with error bars depicting SEM. Statistical significance was determined using two tailed unpaired student’s t test. *=p<0.05, **=p<0.01, ***=p<0.001, ****=p<0.0001.

Because our lists of alternatively spliced genes in resting and *Salmonella*-infected MΦs did not completely overlap, we hypothesized that SRSF7 mediates distinct alternative splicing events in the two MΦ states. To begin to test this, we prioritized a set of genes with the most significant SRSF7-dependent AS changes (>20% PSI; confidence threshold 0.95) in each condition (UN and +SAL) (**Fig. S1A**). Manual annotation of these 36 genes did not reveal significant enrichment for any functional category and none of the top alternative splicing events occurred in genes with characterized roles in the MΦ innate immune response (**Fig. S1A**).

We first looked at SRSF7-dependent exon 6 skipping in *Dcun1d2* (Ensembl Transcript ID: ENSMUST00000045366.10), a gene associated with Cullin neddylation (Kajigaya et al., 2016). MAJIQ measured significant enrichment of this exon skipping event in both untreated and *Salmonella*-infected *Srsf7* knockdown MΦs (ΔPSI 39.2% in UN and 43.4% in +SAL) (**Fig. 1F-G**). We confirmed this via semi-quantitative RT-PCR using RNA isolated from uninfected MΦs and MΦs treated with lipopolysaccharide (LPS). LPS activates TLR4 in MΦs in a manner analogous to *Salmonella* infection but provides tighter kinetic control experimentally. Consistent with our MAJIQ analysis, we measured significantly higher levels of *Dcun1d2* exon 6 skipping in *Srsf7* knockdown MΦs in both conditions (**Fig. 1H-I**).

Next, we looked at an alternative splicing event computationally predicted to be “activation-dependent” (i.e. only detected in +SAL MΦs) (**Fig. 1D**). In +SAL samples, MAJIQ quantified SRSF7-dependent alternative 3’SS usage in exon 3 of the *Mphosph9* gene (Ensembl Transcript ID: ENSMUST00000184951.8), which encodes an M-phase phosphoprotein (*Srsf7* knockdown #2 (ΔPSI −35.7%) (**Fig. 1J**)). Again, by RT-PCR, we confirmed that this event occurs preferentially in *Srsf7* knockdown MΦs (**Fig. 1K**). Curiously however, we also measured preferential alternative 3’SS usage in untreated *Srsf7* knockdown MΦs (**Fig. 1L**), even though this event was not computationally identified by RNA-seq/MAJIQ. Similarly, SRSF7-dependent inclusion of exon 8 in *Gripap1* (Ensembl Transcript ID ENSMUST00000065932.14) (**Fig. S1B**), and inclusion of exon 35 in *Dop1a* (Ensembl Transcript ID ENSMUST00000190957.7) (**Fig. S1D**) only met our threshold in UN and +SAL conditions, respectively, but could be detected in both macrophage states by RT-PCR (**Fig. S1C, S1E**). These data argue that although SRSF7 is important for making alternative splicing decisions in RAW MΦs, it largely mediates the same events in the presence and absence of LPS. Based on the nature of the genes whose alternative splicing relies on SRSF7 and the lack of specificity for alternative splicing in resting vs. activated MΦs, we can conclude that SRSF7 does not dramatically influence the macrophage innate immune transcriptome at the level of splicing.

### SRSF7 is a positive regulator of interferon stimulated genes (ISGs) in macrophages

In addition to the alternative splicing events reported in Fig. 1, we also measured hundreds of differentially expressed genes in resting and *Salmonella*-infected *Srsf7* knockdown RAW MΦs compared to controls (**Supplementary Table 2**). In resting MΦs, *Srsf7* knockdown impacted expression of many housekeeping genes in resting MΦs, particularly in pathways related to translation regulation (EIF2 signaling and Regulation of eIF4 and p70S6K signaling) and phagosome maturation (**Fig. S2A**). Interestingly, in resting MΦs, we also noted enrichment for SRSF7-dependent genes in pathways like “Communication between innate and adaptive immune cells” (**Fig. S2A**), suggesting that loss of SRSF7 dysregulates innate immune gene expression even in the absence of a stimulus.

Enrichment for pathways related to innate immunity was also evident in *Salmonella*-infected macrophages, where knockdown of *Srsf7* dysregulated genes in pathways related to “role of pattern recognition receptors in recognition of bacteria and viruses,” “interferon signaling,” “role of PKR in interferon induction and antiviral response,” and “activation of IRF by cytosolic pattern recognition receptors” (**Fig. S2B**). Many of these categories contain genes related to the type I interferon (IFN) response, a ubiquitous antiviral response characterized by expression of interferon alpha and beta and interferon stimulated genes (ISGs) (Schoggins, 2019; Schoggins and Rice, 2011). In MΦs, the type I IFN response is activated by the transcription factors IRF3 and IRF7 in response to several immune agonists including LPS/*Salmonella* infection, cytosolic dsDNA, and cytosolic dsRNA (Honda and Taniguchi, 2006; Jefferies, 2019; Schoggins et al., 2011).

For ISGs that are appreciably expressed in resting MΦs (Barrat et al., 2019; Gough et al., 2012), e.g. *Apol9a*, *Apol9b*, *Ifit3*, and *Ifi202b*, *Srsf7* knockdown decreased their abundance in both untreated and +*Salmonella* conditions (**Fig. 2A**; bottom left quadrant). Genes like *Mx1, Zbp1,* and *Ifi209*, which are virtually undetectable in resting cells, only displayed SRSF7-dependence in our *Salmonella*-infected samples (**Fig. 2A**; lower y-axis). Visualizing RNA reads from *Srsf7* knockdown versus control MΦs for two such ISGs, *Zbp1* and *Ifit3*, on the Integrated Genomics Viewer, revealed no changes in intronic reads (**Fig. 2B-C**). This was consistent with our MAJIQ analysis, which did not detect many statistically significant splicing changes in genes annotated as ISGs in *Srsf7* knockdown RAW MΦs (**Fig. 1E, Supplemental Table 1**).

**Figure 2:**
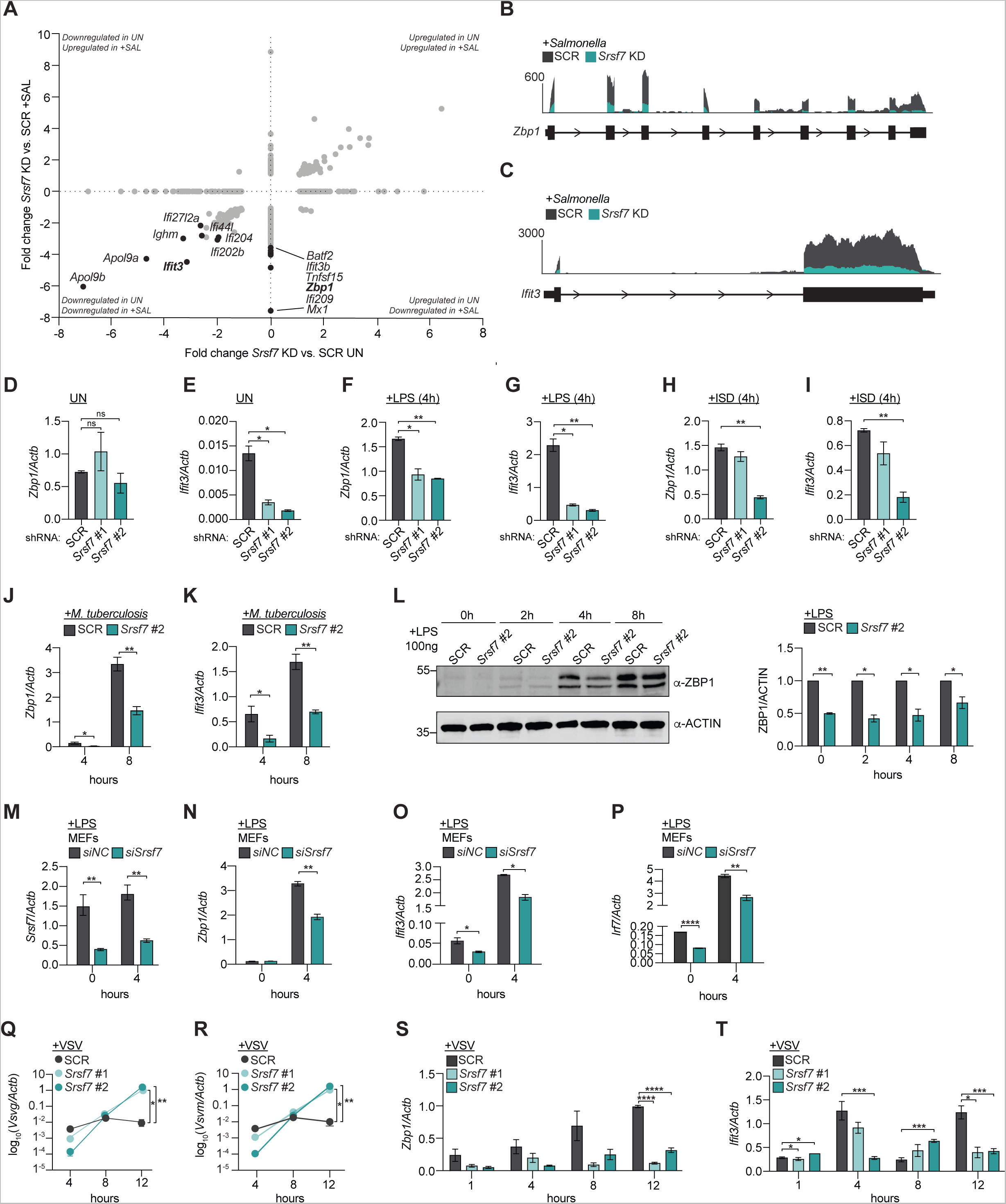
SRSF7 promotes the type I interferon response. **A.** Scatterplot of genes differentially expressed in *Srsf7* KD RAW MΦs relative to SCR in each of the conditions queried (X axis = UN; Y axis = +SAL). Select interferon stimulated genes depicted in black. **B.** Integrated Genomics Viewer track of *Zbp1* SCR (grey) and *Srsf7* KD (teal) reads from *Salmonella*-infected cells. **C.** As in B but for *Ifit3.* **D.** RT-qPCR of *Zbp1* in SCR, *Srsf7* KD#1, and *Srsf7* KD#2 RAW MΦs at baseline (UN). **E.** As in D but for *Ifit3.* **F.** RT-qPCR of *Zbp1* in SCR, *Srsf7* KD#1, and *Srsf7* KD#2 RAW MΦs 4h post-stimulation with 100 ng/ml LPS. **G.** As in F but for *Ifit3.* **H.** RT-qPCR of *Zbp1* in SCR, *Srsf7* KD#1, and *Srsf7* KD#2 RAW MΦs 4h post-transfection with 1 µg/ml ISD. **I.** As in F but for *Ifit3.* **J.** RT-qPCR of *Zbp1* in SCR and *Srsf7* KD #2 RAW MΦs at 4 and 8h post-infection with *M. tuberculosis* (Erdman strain; MOI=5). **K.** As in J but for *Ifit3.* **L.** Immunoblot of ZBP1 protein levels in SCR and *Srsf7* KD #2 RAW MΦs at baseline, 4, and 8h, post-stimulation with 100 ng/ml LPS. Right, quantification. **M.** RT-qPCR of *Srsf7* in primary MEFs at baseline or 4h post stimulation with 100ng LPS. **N.** As in M but for *Zbp1*. **O.** As in M but for *Ifit3*. **P.** As in M but for *Irf7*. **Q.** RT-qPCR of *Vsvg* in SCR, *Srsf7* KD #1, and *Srsf7* KD #2 RAW MΦs at 4, 8, 12h post-infection with VSV (MOI=0.1). **R.** As in M but for *Vsvm.* **S.** RT-qPCR of *Zbp1* in SCR, *Srsf7* KD #1, *Srsf7* KD #2 RAW MΦs at 1, 4, 8, 12h post-infection with VSV (MOI=0.1). **T.** As in S but for *Ifit3.* Data are expressed as a mean of three biological replicates with error bars depicting SEM. Statistical significance was determined using two tailed unpaired student’s t test. *=p<0.05, **=p<0.01, ***=p<0.001, ****=p<0.0001.

**Figure S2:**
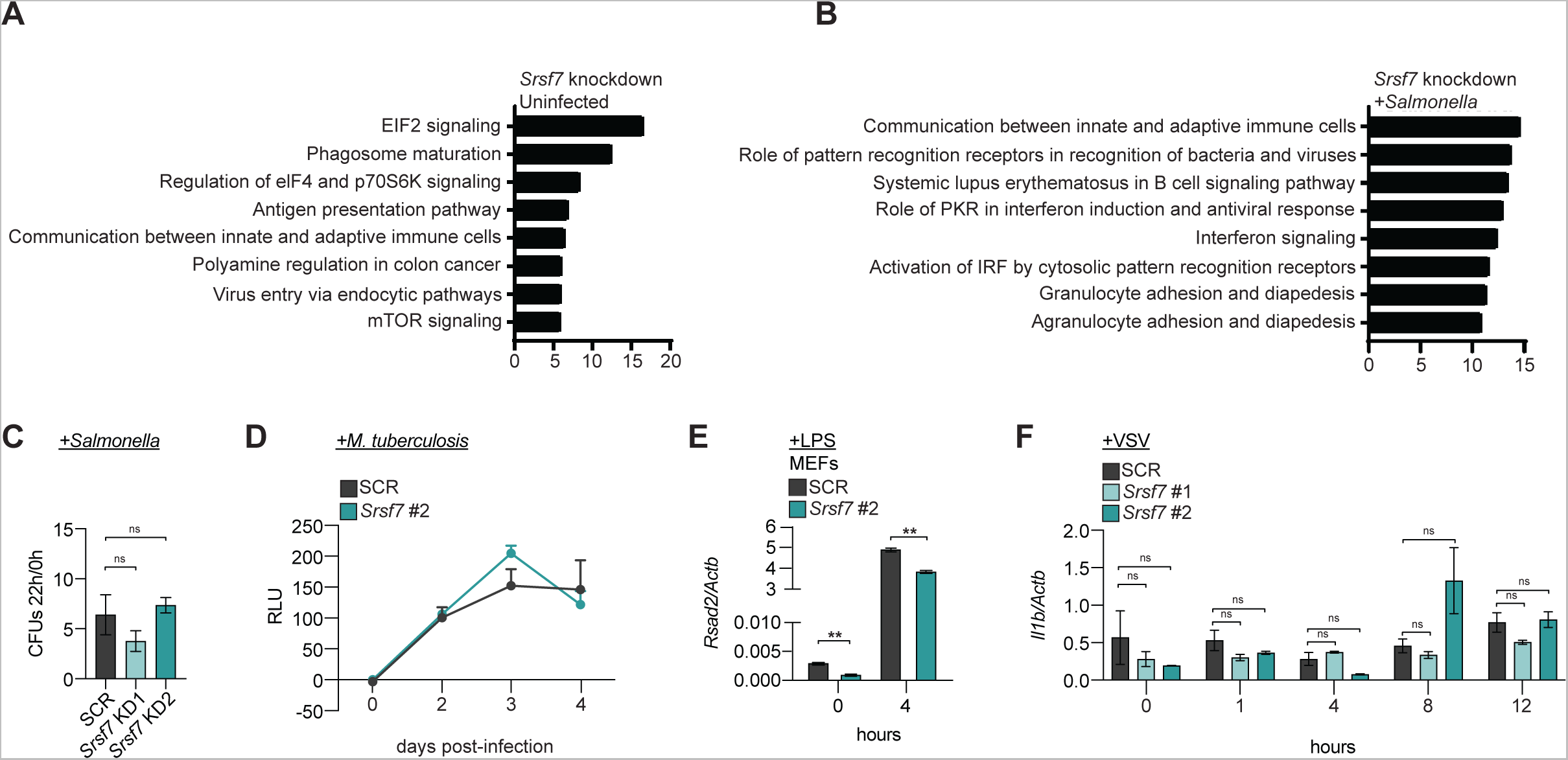
Supplemental data for Figure 2. **A.** Ingenuity Pathway Analysis (IPA; Qiagen) of pathways enriched for differentially expressed genes in *Srsf7* KD RAW MΦs (UN). **B.** As in A for *Srsf7* KD RAW MΦs +SAL. **C.** Infection of SCR and *Srsf7* KD RAW MΦs with *Salmonella* enterica (MOI=1) in non-SPI-1 inducing conditions. Fold-replication represented as 22h over 2h post-infection. **D.** Mtb luxBCADE growth in SCR or *Srsf7* KD RAW MΦs measured by relative light units (RLUs) over a 4-day time course (MOI=1). **E.** RT-qPCR of *Rsad2* in siRNA *Srsf7-* or siControl-transfected primary MEFs at baseline or 4h post stimulation with 100ng/ml of LPS. **F.** RT-qPCR of *Il1b* in SCR or *Srsf7* KD RAW MΦs over a time course of VSV infection. Data are expressed as a mean of three or more biological replicates with error bars depicting SEM. Statistical significance was determined using two tailed unpaired student’s t test. *=p<0.05, **=p<0.01.

Failure to phagocytose *Salmonella* or defects in *Salmonella* replication could result in lower ISG expression in *Srsf7* knockdown cells, but enumeration of colony forming units (CFUs) did not reveal any differences in bacterial burdens between SCR and *Srsf7* knockdown RAW MΦs over a 22h infection time-course (**Fig. S2C**). To determine if SRSF7’s ability to promote ISG expression was unique to *Salmonella* infection, we measured ISG expression in SCR and *Srsf7* knockdown MΦs at 4h post-treatment with LPS (100 ng/ml), after dsDNA transfection, which activates type I IFN gene expression downstream of cGAS (1 μg ISD), and after treatment with recombinant IFN-β which activates ISG expression in MΦs by binding to IFNAR and engaging JAK/STAT cascades (200 IU, 4h). In all cases, loss of *Srsf7* led to impaired activation of *Zbp1* and *Ifit3* expression (**Fig. 2D-I**). We also measured lower ISG expression in *Srsf7* knockdown MΦs during infection with the intracellular bacterial pathogen, *Mycobacterium tuberculosis* (MOI = 5) (**Fig. 2J-K**), which elicits *Ifnb1* and ISG expression via the cGAS/STING pathway (Watson et al., 2015). Again, this defect in ISG expression was independent of bacterial internalization/replication (**Fig. S2D**). Over an 8h time-course of LPS-treatment, lower *Zbp1* transcript levels correlated with lower protein accumulation (**Fig. 2L**), suggesting that failure to fully upregulate ISG expression in *Srsf7* knockdown MΦs is likely to functionally impact the innate immune response to infection. This phenotype was not unique to RAW MΦs, as siRNA knockdown of SRSF7 in primary mouse embryonic fibroblasts (MEFs)(**Fig. 2M**), a cell type commonly used for studies of type I interferon signaling, also resulted in low basal ISG expression and a failure to fully induce ISGs upon LPS stimulation (**Fig. 2N-P and S2E**).

To test whether SRSF7 is a *bona fide* player in activating antimicrobial immunity in MΦs, we infected SCR and *Srsf7* knockdown MΦs with vesicular stomatitis virus (VSV), a negative-strand RNA virus that stimulates type I IFN responses via TLR3 and RIG-I and is extremely sensitive to ISG expression (Gao et al., 2021; Wagner et al., 2021). We measured a dramatic increase in VSV replication in *Srsf7* knockdown MΦs, measured by RT-qPCR of two VSV genes, *Vsvg* and *Vsvm* (**Fig. 2Q-R**), that was concomitant with depressed ISG transcript induction over a 12-hour time course of VSV infection (MOI = 0.1) (**Fig. 2S-T**). We did not detect SRSF7-dependent differences in *Il1b* expression in response to VSV infection, arguing against a role for SRSF7 in controlling innate gene expression downstream of NFκB (**Fig. S2F**). Together, these data identify SRSF7 as a positive regulator of the type I IFN response in MΦs and suggest that the mechanism of SRSF7-dependent ISG expression is shared by several pathogen sensing cascades (i.e. occurs at one or more nodes where the TLR4, cGAS, and RIG-I cascades converge).

### SRSF7 drives ISG expression by promoting expression of the transcription factor IRF7

Given that loss of SRSF7 results in a failure to fully induce many ISGs, we hypothesized that SRSF7 may act directly on one or more critical innate signaling proteins or transcription factors. To pinpoint where in the type I IFN signaling response SRSF7 acts, we treated *Srsf7* knockdown or SCR RAW MΦs with LPS (100 ng/ml) and measured secretion of type I interferon (IFN-βα) over a 6h time-course via an ISRE-luciferase reporter cell line (Hoffpauir et al., 2020; Wagner et al., 2022). *Srsf7* knockdown MΦs produced significantly less IFN-βα compared to SCR controls at all time points measured (**Fig. 3A**). We next asked if impaired ISG expression in *Srsf7* knockdown MΦs could be rescued if *Srsf7* knockdown and SCR control cells received equal amounts of recombinant IFN-β. We found that even post treatment of IFN-β, *Srsf7* knockdown MΦs fail to express ISGs like *Ifi202b* and *Zbp1* to the same level as SCR controls, at both the transcript (**Fig. 3B-C**) and protein level (**Fig. 3D**). These results suggest that loss of SRSF7 impacts some aspect of the type I IFN response that is engaged upstream and downstream of IFNAR and that loss of IFN-β production is not the sole driver of ISG defects in *Srsf7* knockdown MΦs.

**Figure 3:**
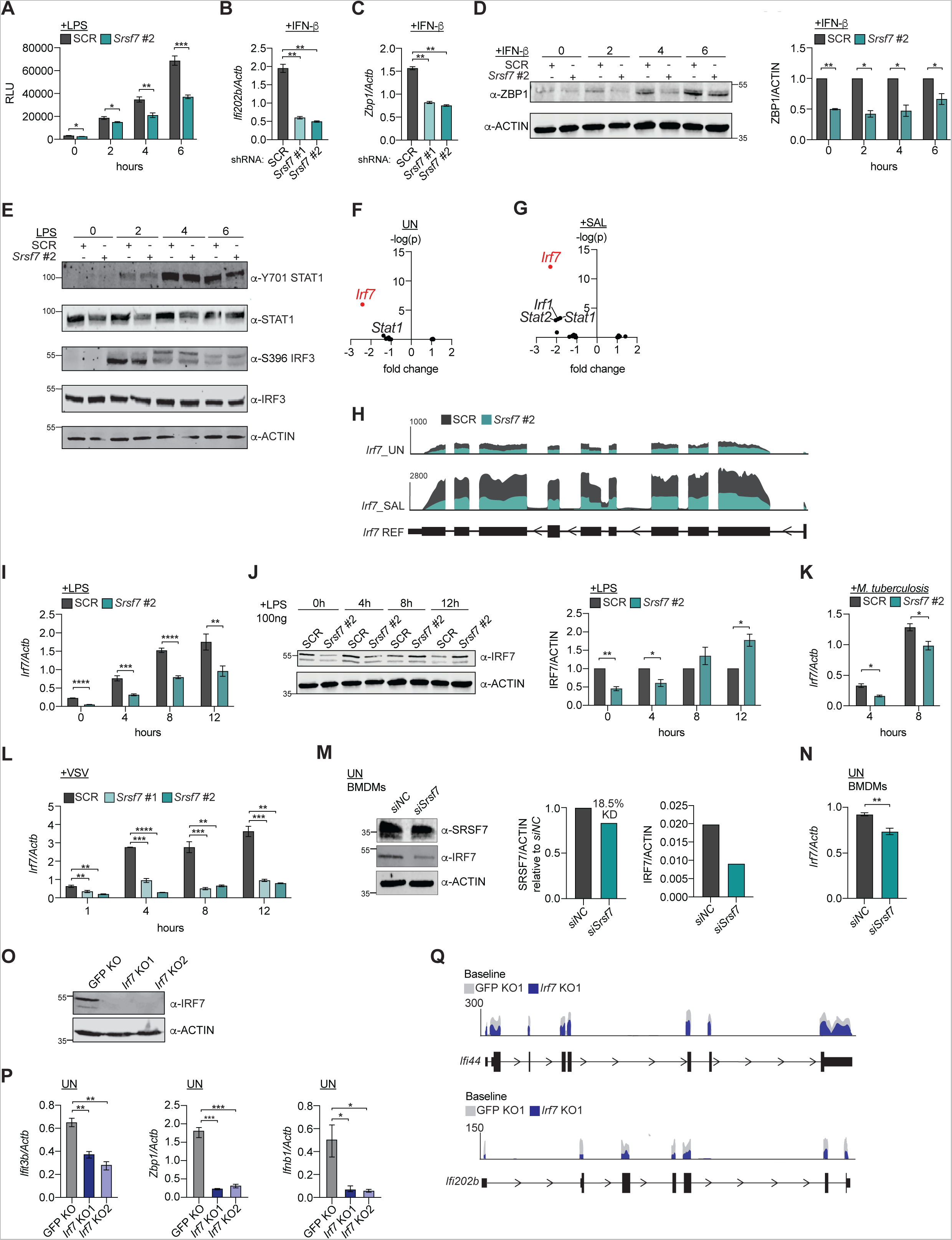
SRSF7 is required for IRF7 accumulation in macrophages. **A.** Secreted IFNα/β in SCR and *Srsf7* KD#2 RAW MΦs, as a measure of luciferase expressed by ISRE reporter cells. **B.** RT-qPCR of *Ifi202b* in SCR, *Srsf7* KD#1 and *Srsf7* KD#2 in RAW MΦs 4h post-stimulation with 200 IU IFN-β. **C.** As in B for *Zbp1.* **D.** Immunoblot of ZBP1 and ACTIN over a time course of IFN-β (200 IU) treatment in SCR and *Srsf7* KD#2 RAW MΦs. Quantification, right. **E.** Immunoblot of Y701 STAT1, STAT1, S396 IRF3, IRF3, and ACTIN in SCR and *Srsf7* KD RAW MΦs over a time course of LPS treatment (100 ng/ml). **F.** Differential expression of transcription factor genes in untreated *Srsf7* knockdown vs. SCR RAW MΦs as fold change and −log(p-value). Gene of interest (*Irf7*) highlighted in red. **G.** As in F but for +SAL samples. **H.** Integrated Genomics Viewer with *Irf7* reads from SCR (grey) and *Srsf7* KD#2 (teal) RAW MΦs, untreated (top) and +*Salmonella* (bottom). **I.** RT-qPCR of *Irf7* over a time course of LPS stimulation (100 ng/ml) in SCR and *Srsf7* KD RAW MΦs. **J.** Immunoblot of IRF7 and ACTIN protein levels over a time course of LPS stimulation (100 ng/ml) in SCR and *Srsf7* KD RAW MΦs. Quantification, right. **K.** RT-qPCR of *Irf7* over a time course of Mtb infection (Erdman strain; MOI=5) in SCR and *Srsf7* KD#2 MΦs. **L.** RT-qPCR of *Irf7* over a time course of VSV infection (MOI=0.1) in SCR, *Srsf7* KD#1, *Srsf7* KD#2 RAW MΦs. **M.** Immunoblot of SRSF7, IRF7, and ACTIN in siControl- or si*Srsf7*-transfected BMDMs at baseline. Quantifications, right. **N.** RT-qPCR of *Irf7* in siControl or si*Srsf7* BMDMs at baseline. **O.** Immunoblot of IRF7 protein levels in GFP gRNA, *Irf7* KO1 and *Irf7* KO2 RAW MΦs at baseline. ACTIN used as loading control. **P.** RT-qPCR of *Ifit3b, Zbp1,* or *Ifnb1* at baseline in *Irf7* KO1 (blue) vs. GFP gRNA control (grey) RAW MΦs. **Q.** Integrated Genomics Viewer with *Ifi44* and *Ifi202b* reads from GFP gRNA (grey) or *Irf7* KO (blue) RAW MΦs at baseline (UN). Data expressed as a mean of three biological replicates with error bars depicting SEM unless otherwise noted. Statistical significance was determined using two tailed unpaired student’s t test. *=p<0.05, **=p<0.01, ***=p<0.001, ****=p<0.0001.

We then asked whether altered signaling downstream of TLR4 could explain defective ISG expression in *Srsf7* knockdown RAW MΦs. We treated *Srsf7* knockdown and SCR RAW MΦs with LPS and measured phosphorylation of the transcription factor IRF3 (activated downstream of TLR4-TRIF signaling) and STAT1 (activated downstream of IFNAR) over a 6h time course. We did not measure significant changes in phosphorylation of IRF3 at S396 or phosphorylation of STAT1 at Y701 (**Fig. 3E**; quantification in **Fig. S3A**), effectively ruling out defective signal transduction as the culprit for dampened ISG expression in *Srsf7* knockdown MΦs.

Next, we entertained the possibility that SRSF7 is required for the expression of one or more transcription factors needed to activate the type I IFN response downstream of pattern recognition receptor and/or IFNAR engagement. To test this, we generated a list of transcription factors associated with macrophage innate immunity in the STAT, IRF, and NFκB families, and turned back to our RNA-seq data to see how expression of each factor was impacted by loss of SRSF7. While expression of most transcription factors was unaffected, we observed a significant defect in *Irf7* expression in *Srsf7* knockdown RAW MΦs at baseline and after *Salmonella* infection (**Fig. 3F-G**). Visualization of *Irf7* gene expression data on the IGV highlights fewer *Irf7* exonic reads in *Srsf7* knockdown (teal) compared to SCR (grey) RAW MΦs in both conditions (**Fig. 3H**). We did not detect an increase in intron reads in *Srsf7* knockdown MΦs nor did our MAJIQ analysis identify SRSF7-dependent *Irf7* alternative splicing events (**Fig. 1D**), arguing against a role for SRSF7 in regulating *Irf7* transcript abundance via splicing.

Initially discovered in the context of Epstein Barr virus, IRF7 (interferon regulatory factor 7) is a master regulator of antiviral type I IFN responses (Honda et al., 2005). Importantly, IRF7 is itself an ISG whose expression is induced downstream of IFNAR engagement. Although basal levels of IRF7 vary between immune cell types, we know that dendritic cells are heavily reliant upon IRF7 at baseline and several studies have implicated IRF7 in priming early responses in macrophages downstream of LPS (Fitzgerald et al., 2003; Gu et al., 2022; Jefferies, 2019; Sin et al., 2020). We can readily measure IRF7 transcript and protein in resting wild-type RAW MΦs (**Fig. 3I-J**) and in wild-type immortalized BMDMs (**Fig. S3B**), confirming their utility for studies of IRF7 in resting and activated states. IRF7 protein levels are low in WT resting BMDMs and are upregulated in response to LPS (**Fig. S3C**).

To better appreciate the kinetics of SRSF7-dependent control of *Irf7* expression, we treated *Srsf7* knockdown and SCR control RAW MΦs with LPS and measured *Irf7* transcript and protein over a 12h time course. Levels of *Irf7* mRNA were significantly lower in *Srsf7* knockdown RAW MΦs at every time-point queried (**Fig. 3I**). IRF7 protein levels were lower in *Srsf7* knockdown RAW MΦs at 0 and 4h but appeared to rebound at later time points (8 and 12h post-LPS) (**Fig. 3J**), consistent with earlier data suggesting that IRF7 protein can be stabilization during the innate response to infection (Ning et al., 2011). We also measured significantly less *Irf7* transcript in *Srsf7* knockdown RAW MΦs in response to *M. tuberculosis* (**Fig. 3K**) and VSV infection (**Fig. 3L**). Although we were only able to achieve modest knockdown of SRSF7 in BMDMs transfected with an siRNA directed against *Srsf7* (alongside a non-targeting control; siNC) (**Fig. 3M**), we can detect less IRF7 protein (**Fig. 3M**) and transcript (**Fig. 3N**) in these primary MΦs as well. Together, our data demonstrate a previously unappreciated role for SRSF7 in controlling *Irf7* transcript and protein levels in macrophage cell lines, primary macrophages, (**Fig. 3I-J** and **3M-N**) and MEFs (**Fig. 2P**).

**Figure S3:**
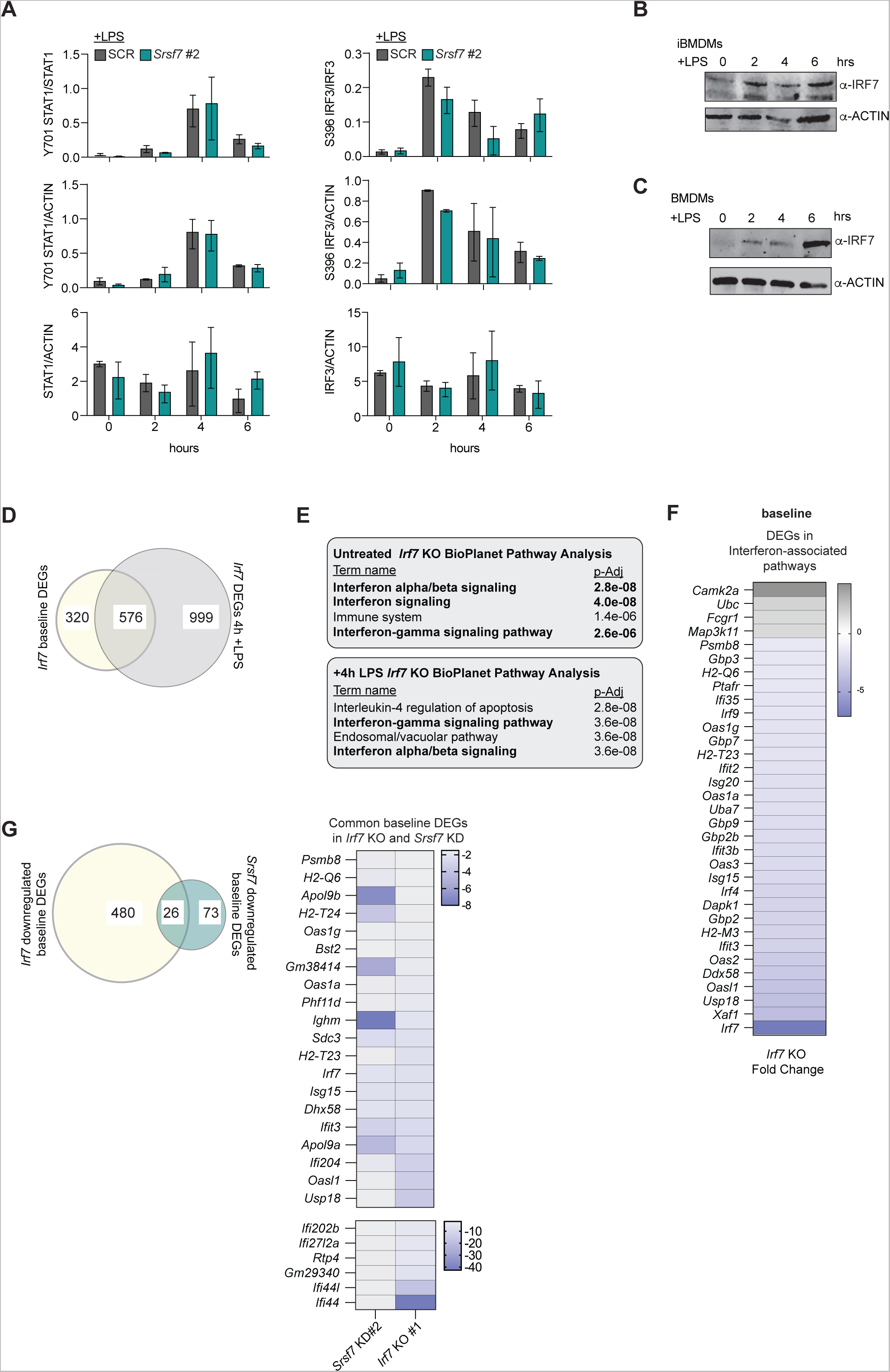
Supplemental data for Figure 3. **A.** Quantification of Figure 3E; n=2 **B.** Immunoblot of IRF7 and ACTIN protein levels over a time course of LPS stimulation (100 ng/ml) in WT iBMDMs. **C.** As in B but in WT BMDMs. **D.** Venn diagram depicting differential gene expression of *Irf7* KO1 at baseline vs. 4h +LPS (100ng/ml). Cut off: fold change < −1.5 and p-value <0.05. **E.** Top: BioPlanet Pathway Analysis of untreated *Irf7* KO gene expression. Bottom: BioPlanet Pathway Analysis of 4h +LPS *Irf7* KO gene expression. **F.** Heatmap depicting *Irf7* KO baseline gene expression from BioPlanet pathway analysis categories: “Interferon alpha/beta signaling,” “Interferon signaling,” and “Interferon-gamma signaling pathway.” Duplicate terms removed. **G.** Left: Venn diagram depicting differentially expressed genes (DEGs) in *Irf7* KO1 (compared to GFP gRNA) and *Srsf7* KD (compared to SCR) RAW MΦs at baseline. Right: Heatmap of 26 genes in common between *Irf7* KO1 and *Srsf7* KD at baseline. Data are expressed as a mean of three biological replicates with error bars depicting SEM unless otherwise noted. Statistical significance was determined using two tailed unpaired student’s t test.

### IRF7 controls basal and induced expression of ISGs in RAW MΦs

Our findings argue that loss of ISG expression in *Srsf7* knockdown cells stems primarily from dampened levels of IRF7. However, as most literature points to a role for IRF7 in activated immune cells, we were initially cautious to attribute basal ISG phenotypes in *Srsf7* knockdown MΦs to IRF7. To better understand how IRF7 controls type I IFN responses in resting and activated RAW MΦs, we generated *Irf7* KO RAW MΦs using CRISPR-Cas9 editing with a gRNA designed to target *Irf7* exon 3, and negative control cells expressing a GFP-specific (non-targeting) gRNA. Two clonally expanded *Irf7* KO cell lines and a GFP gRNA control cell line were selected, and loss of IRF7 protein was confirmed by immunoblot (**Fig. 3O**). Total RNA from untreated *Irf7* KO #1 and control cell lines, as well as from cells stimulated with LPS for 4h (100 ng/ml) were sent for RNA-seq analysis. We measured differential expression of 896 transcripts (fold-change < −1.5; p<0.05) in *Irf7* KO vs. control cells at baseline and 1575 genes were differentially expressed in *Irf7* KO MΦs at 4h post-LPS treatment (**Supplemental Table 3** and **Fig. S3D**). Pathway analysis (BioPlanet Pathway Enrichment Analysis, NCATS (Huang et al., 2019)) identified a number of type I interferon-related pathways enriched for differentially expressed genes in *Irf7* KO cells at baseline and at 4h post-LPS treatment (**Fig. S3E**). Consistent with our hypothesis that SRSF7 controls ISG expression in an IRF7-dependent fashion, we found that ISG expression was dramatically dampened in *Irf7* KO RAW MΦs at rest and after LPS stimulation (**Figs. 3P-Q**, **S3F**, and **Supplemental Table 3**) and that differentially expressed ISGs were largely shared between *Srsf7* KD and *Irf7* KO cells (**Fig. S3G**). Together, these data provide strong evidence for IRF7 maintaining homeostatic expression of type I IFN and ISGs in macrophages and suggest that SRSF7 contributes to this circuit by mediating IRF7 expression.

### SRSF7 overexpression is sufficient to drive IRF7 and ISG expression

We hypothesized that if SRSF7 can directly activate *Irf7* transcription, then overexpression of SRSF7 should be sufficient to drive *Irf7* expression. To test this model, we generated doxycycline-inducible RAW MΦ cell lines that express 3xFLAG-SRSF7 or 3xFLAG-GFP. To ensure comparable levels of 3xFLAG-GFP and 3xFLAG-SRSF7 protein expression and to best mimic endogenous levels of SRSF7, we tested and optimized treatment times, deciding on 2h +DOX for 3xFLAG-GFP and 22h +DOX for 3xFLAG-SRSF7 (**Fig. 4A**). Notably, we found that expression of SRSF7 alone was sufficient to increase the abundance of *Irf7* transcript (**Fig. 4B**), IRF7 protein levels (**Fig. 4C**), and ISG transcripts (**Fig. 4D**) in otherwise unstimulated RAW MΦs. Doxycycline-inducible expression of 3xFLAG-SRSF7 was also sufficient to drive expression of additional genes in the IRF7 baseline transcriptional regulon (*Xaf*, *Irak2*, *Ccl2*, and *Isg20* (**Figs. 4E and S3G**)), lending further support to our hypothesis that SRSF7 controls ISG expression at the level of IRF7. We observed a similar phenotype in RAW MΦ cell lines that stably express 3xFLAG-SRSF7 compared to 3xFLAG-GFP controls (**Fig. 4F**). While overexpression of 3xFLAG-SRSF7 had a less pronounced impact on IRF7 transcript and protein expression after LPS stimulation (**Fig. S4A-B**), we measured a dramatic increase in *Irf7* and *Ifit3* transcript levels in 3xFLAG-SRSF7-expressing stable cells upon infection with *M. tuberculosis* (**Fig. 4G**). Because expression of interferon stimulated genes has been associated with cell-intrinsic antibacterial control mechanisms and cell death pathways that impact bacterial spread (Banks et al., 2019; Hoffpauir et al., 2020; MacMicking et al., 1997; MacMicking et al., 2003), we also asked whether overexpression of SRSF7 affected *M. tuberculosis* replication. Briefly, we infected cells with Mtb *luxBCADE* and followed bacterial replication as a measure of relative light units (as in (Bell et al., 2021; Wagner et al., 2022)). We measured a significant increase in RLUs at Day 3 post-infection (**Fig. 4H**) suggesting that SRSF7 expression may help control *M. tuberculosis* replication and/or spread. Together, these data suggest a direct role for SRSF7 in activating expression of *Irf7* and promoting type I interferon responses in MΦs.

**Figure 4:**
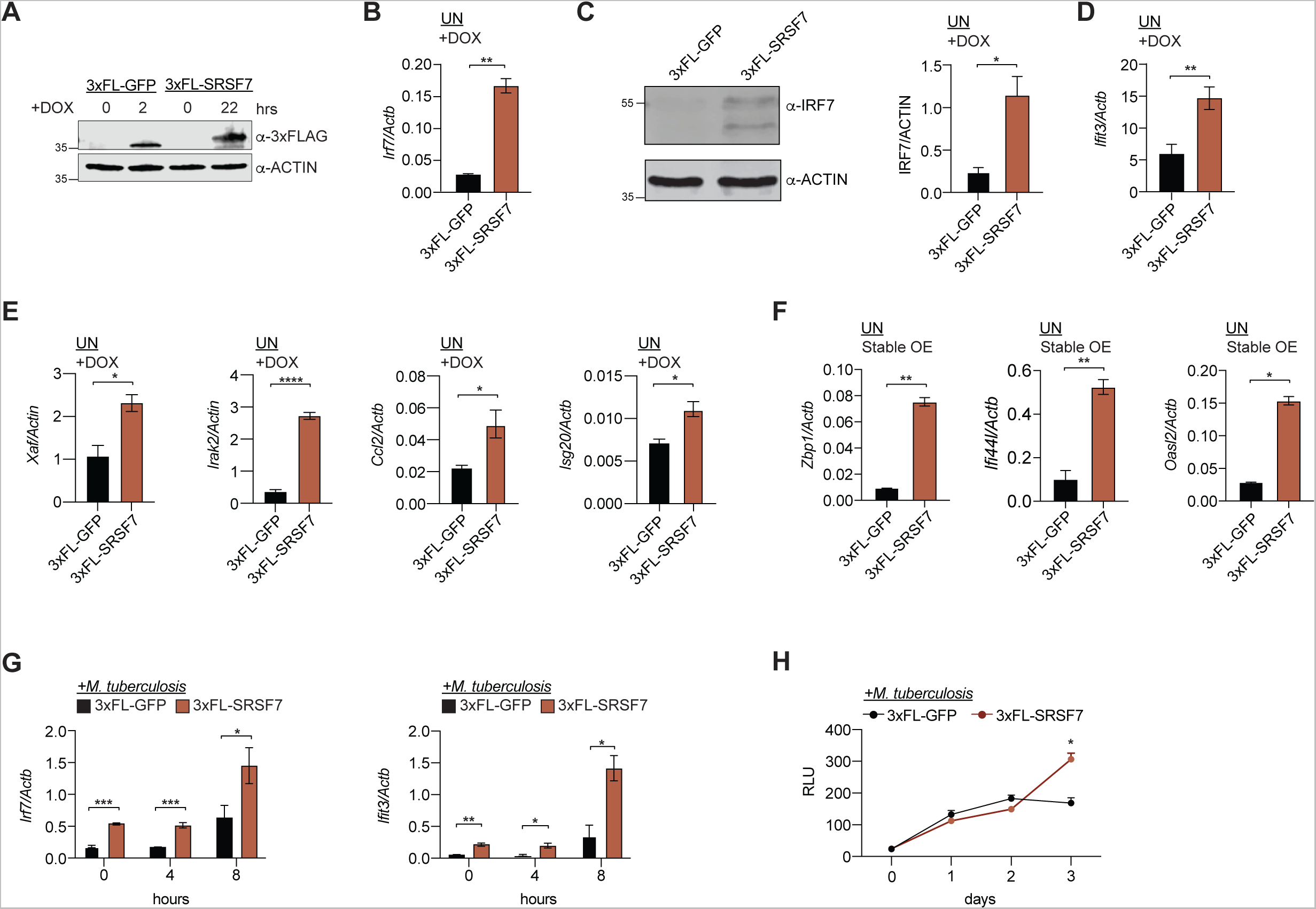
SRSF7 overexpression is sufficient to drive basal expression of IRF7 and ISGs. **A.** Immunoblot of 3xFLAG-GFP or 3xFLAG-SRSF7 tetracycline inducible RAW MΦs treated with doxycycline (DOX) (1 µg/ml). 3xFLAG-GFP MΦs received DOX for 2h and 3xFLAG-SRSF7 MΦs received DOX for 22h. ACTIN shown as loading control. **B.** RT-qPCR of *Irf7* in 3xFLAG-GFP (2h DOX treatment) and 3xFLAG-SRSF7 (22h DOX treatment) inducible RAW MΦs at baseline. **C.** Immunoblot of IRF7 in 3xFLAG-GFP (2h DOX treatment) and 3xFLAG-SRSF7 (22h dox treatment) inducible RAW MΦs at baseline. Quantification right. **D.** As in B but for *Ifit3*. **E.** As in B but for *Xaf, Irak2, Ccl2,* and *Isg20*. **F.** RT-qPCR of *Zbp1*, *Ifi44*, or *Oasl2* in 3xFLAG-GFP and 3xFLAG-SRSF7 stable overexpression RAW MΦs at baseline. **G.** RT-qPCR of *Irf7* or *Ifit3* at 0, 4, and 8h post-infection with Mtb (Erdman; MOI=5). **H.** Mtb *luxBCADE* replication in 3xFLAG-GFP and 3xFL-SRSF7 doxycycline-inducible RAW MΦs measured by relative light units (RLUs) over time course (Erdman; MOI=1). Data are expressed as a mean of three biological replicates with error bars depicting SEM. Statistical significance was determined using two tailed unpaired student’s t test. *=p<0.05, **=p<0.01, ***=p<0.001.

**Figure S4:**
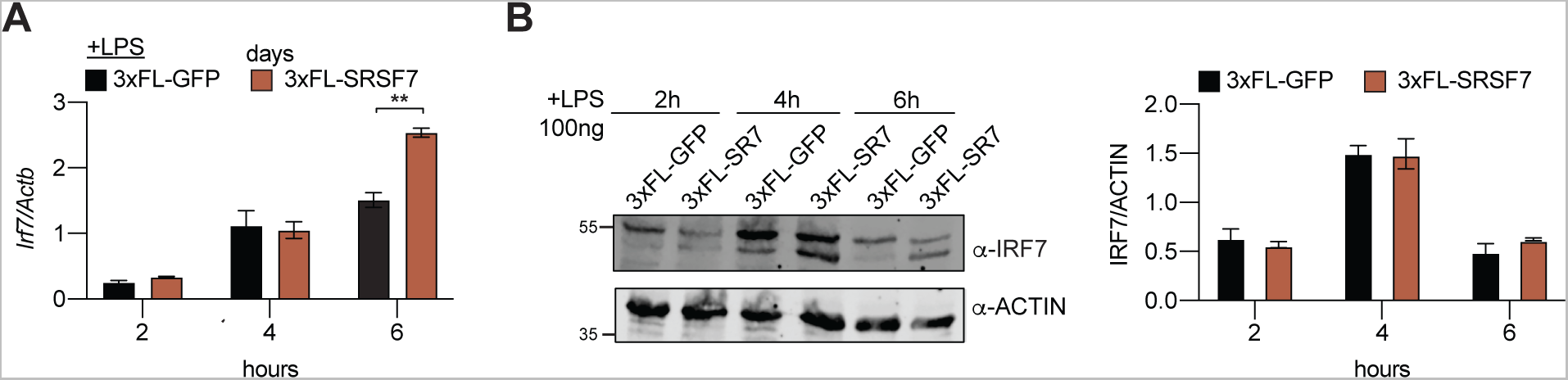
Supplemental data for Figure 4. **A**. RT-qPCR of *Irf7* in 3xFLAG-GFP (2h dox treatment) and 3xFLAG-SRSF7 (22h dox treatment) inducible RAW MΦs over a time course of LPS stimulation (100 ng/ml). **B.** Immunoblot of IRF7 in 3xFLAG-GFP (2h dox treatment) and 3xFLAG-SRSF7 (22h dox treatment) inducible RAW MΦs over a time course of LPS stimulation (100 ng/ml) (quantification right). Data are expressed as a mean of three biological replicates with error bars depicting SEM. Statistical significance was determined using two tailed unpaired student’s t test. *=p<0.05, **=p<0.01, ***=p<0.001.

### SRSF7 promotes STAT1 association and RNAPII elongation at the *Irf7* promoter

Having established that decreased levels of IRF7 is the likely culprit for the type I IFN defect in *Srsf7* knockdown MΦs (**Fig. 3**) and demonstrated that SRSF7 overexpression is sufficient to promote *Irf7* expression (**Fig. 4**), we set out to determine the mechanism though which SRSF7 controls *Irf7* expression. Earlier work has implicated SR proteins in coupling transcription and splicing and have been shown to influence transcription by interacting with histone tails (Loomis et al., 2009; Luco et al., 2011; Luco et al., 2010) or by directly mediating transcriptional elongation via pTEFb (Ji et al., 2013). To begin to investigate a role for SRSF7 in activating *Irf7* transcription, we performed chromatin immunoprecipitation (ChIP)-qPCR for RNA polymerase II (RNAPII) in *Srsf7* knockdown and SCR control MΦs at 4h post-LPS treatment, using primers designed to tile the *Irf7* promoter region (100bp amplicon, 100 bp apart from each other (**Fig. 5A**)) and an antibody that recognizes both phosphorylated and unphosphorylated forms of the C-terminal domain of the largest subunit of RNAPII. We observed dramatic accumulation of RNAPII at/around the *Irf7* transcription start site (TSS) (−400 to +250) in *Srsf7* knockdown MΦs (**Fig. 5B**, teal). This pattern of RNAPII in *Srsf7* knockdown MΦs is suggestive of a pile-up resulting from a failure to convert to a successful elongation complex. We next asked whether a defect in RNAPII clearance could be explained by defects in recruitment or retention of transcription factors like STAT1 to the *Irf7* promoter. Using ChIP-qPCR with an antibody against Y701 STAT1, we measured a near total loss of STAT1 signal in *Srsf7* knockdown MΦs (**Fig. 5C**). An analogous set of ChIP-qPCRs at the promoter of *Ifit3*, another ISG whose expression is SRSF7-dependent, showed no differences in RNAPII or STAT1 occupancy in *Srsf7* knockdown cells (**Figs. 5D-F**). Likewise, we saw no difference in RNAPII occupancy in *Srsf7* knockdown MΦs at the promoter of *Stat1* itself (**Figs. 5G-H**). Consistent with upregulation of *Irf7* transcript and protein levels (**Figs. 4B-C**), overexpression of 3xFLAG-SRSF7 was sufficient to recruit significant levels of STAT1 to the *Irf7* promoter in resting macrophages (**Fig. 5I**). These data support a role for SRSF7 in promoting transcription of *Irf7* and suggest that dampened expression of ISGs like *Ifit3* in *Srsf7* knockdown MΦs occurs indirectly, downstream of *Irf7* transcriptional defects.

**Figure 5:**
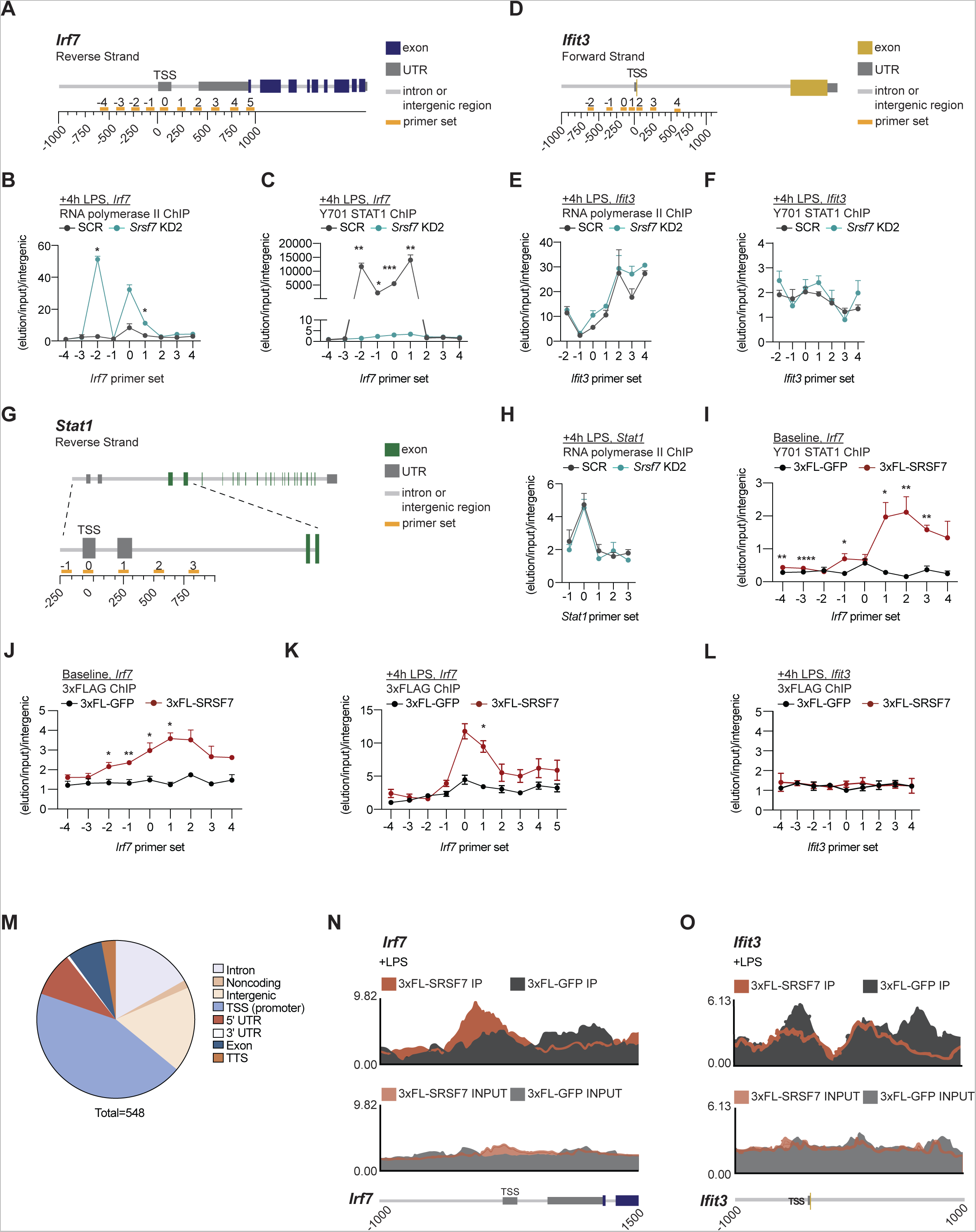
SRSF7 recruitment to the *Irf7* promoter maximizes STAT1 association and promotes RNA polymerase II elongation. **A.** Diagram of the *Irf7* gene with tiling primers designed around the transcriptional start site (orange). **B.** RNA polymerase II ChIP-qPCR (antibody recognizes both modified and unmodified CTD) at the *Irf7* promoter in SCR and *Srsf7* KD RAW MΦs 4h post-LPS stimulation (100ng/ml). Data represented as elution/input normalized to that of an intergenic control primer set. **C.** Y701 STAT1 ChIP-qPCR at the *Irf7* promoter in SCR or *Srsf7* KD RAW MΦs 4h post-LPS stimulation (100 ng/ml). Data represented as elution/input normalized to that of an intergenic control. **D.** Diagram of the *Ifit3* gene with tiling primers designed around the transcriptional start site (orange). **E.** As in B but at the *Ifit3* promoter. **F.** As in C but at the *Ifit3* promoter. **G.** Diagram of the *Stat1* gene with tiling primers designed around the transcriptional start site (orange). **H.** As in B but at the *Stat1* promoter. **I.** Y701 STAT1 ChIP-qPCR at the *Irf7* promoter in 3xFLAG-GFP and 3xFLAG-SRSF7 dox-inducible RAW MΦs at baseline. **J.** 3xFLAG ChIP-qPCR of 3xFLAG-GFP and 3xFLAG-SRSF7 in dox-inducible RAW MΦs at baseline. **K.** 3xFLAG ChIP-qPCR of 3xFLAG-GFP and 3xFLAG-SRSF7 in dox-inducible RAW MΦs at the *Irf7* promoter 4h post LPS stimulation (100 ng/ml). **L.** As in K at the *Ifit3* promoter. **M.** Categorization of unique 3xFLAG ChIP peaks in 3xFL-SRSF7 dox-inducible RAW MΦs 4h post-LPS (100 ng/ml). **N.** Enrichment of 3xFLAG-SRSF7 and 3xFLAG-GFP at the *Irf7* promoter by ChIP-seq: IP (top) and input (bottom). **O.** As in M but at the *Ifit3* promoter. Unless otherwise noted, data are expressed as a mean of three or more biological replicates with error bars depicting SEM. ChIP-qPCRs are shown as the mean of three or more technical replicates with error bars depicting SEM. Statistical significance was determined using two tailed unpaired student’s t test. *=p<0.05, **=p<0.01, ***=p<0.001.

**Figure S5:**
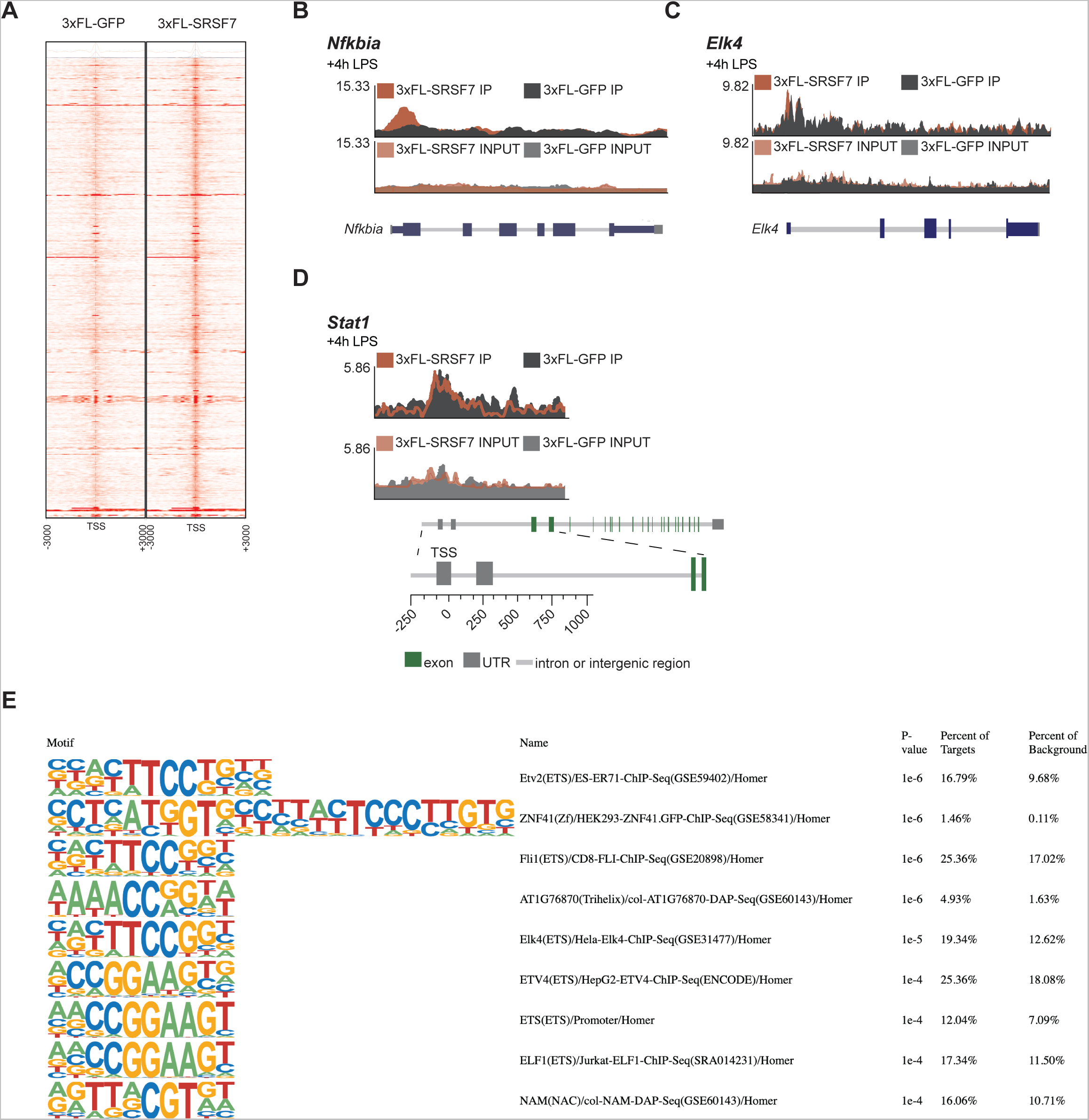
Supplemental data for Figure 5. **A.** Heatmap indicating read coverage density at transcriptional start sites (TSS) in 3xFL-GFP or 3xFL-SRSF7 dox-inducible RAW MΦs, 4h post-LPS treatment. **B.** Enrichment of 3xFLAG-SRSF7 and 3xFLAG-GFP at the *Nfkbia* promoter by ChIP-seq: IP (top) and input (bottom). **C.** Enrichment of 3xFLAG-SRSF7 and 3xFLAG-GFP at the *Elk4* promoter by ChIP-seq: IP (top) and input (bottom). **D.** Enrichment of 3xFLAG-SRSF7 and 3xFLAG-GFP at the *Stat1* promoter by ChIP-seq: IP (top) and input (bottom). **E.** Putative SRSF7 motifs predicted by HOMER from 3xFLAG-SRSF7 ChIP-seq data.

Having linked STAT1 binding and RNAPII elongation at the *Irf7* promoter to SRSF7, we next asked whether SRSF7 carries out this function by directly associating with the *Irf7* genomic locus. Lacking ChIP-grade antibodies for SRSF7, we turned back to our doxycycline-inducible 3xFLAG-SRSF7 and 3xFLAG-GFP cell lines (**Fig. 4A**) and performed anti-FLAG ChIP-qPCR at the *Irf7* promoter using the same primer sets shown in Fig. 6A. Consistent with a direct role for SRSF7 in facilitating STAT1 recruitment and RNAPII elongation at *Irf7*, we observed significant enrichment of 3xFLAG-SRSF7 at the *Irf7* promoter (−500 to 0) in both resting (**Fig. 5J**) and LPS-treated macrophages (4h post-LPS) (**Fig. 5K**) with no SRSF7 enrichment at *Ifit3* (**Fig. 5L**). To determine the extent to which SRSF7 globally associates with promoter regions, we performed ChIP-seq. Briefly, reads were trimmed by Cutadapt and aligned to the *Mus musculus* genome build mm10 using Bowtie. After quality control, peaks were called using MACS2 with input/IgG controls background subtracted. We detected 548 unique peaks for SRSF7 across the genome, with approximately half of these peaks detected at transcription start sites (**Figs. 5M and S5A-C**). By ChIP-seq we again saw clear enrichment of 3xFLAG-SRSF7 at the *Irf7* promoter (**Fig. 5N**), but not at *Ifit3* (**Fig. 5O**) or *Stat1* (**Fig. S5D**). Although SRSF7 peaks were called at several immune associated genes, including *Fos*, *Myd88*, *Ifitm3*, and *Dusp5*, there was no evidence for SRSF7 enrichment at the promoter of any other type I IFN response-associated transcription factors (**Supplemental Table 4**). Taken together, our ChIP-seq data argue against universal association of SRSF7 with gene promoters, as was reported for factors like SRSF1 and SRSF2 (Ji et al., 2013), and instead support a privileged role for SRSF7 in facilitating transcription at specific genes like *Irf7*.

**Figure 6:**
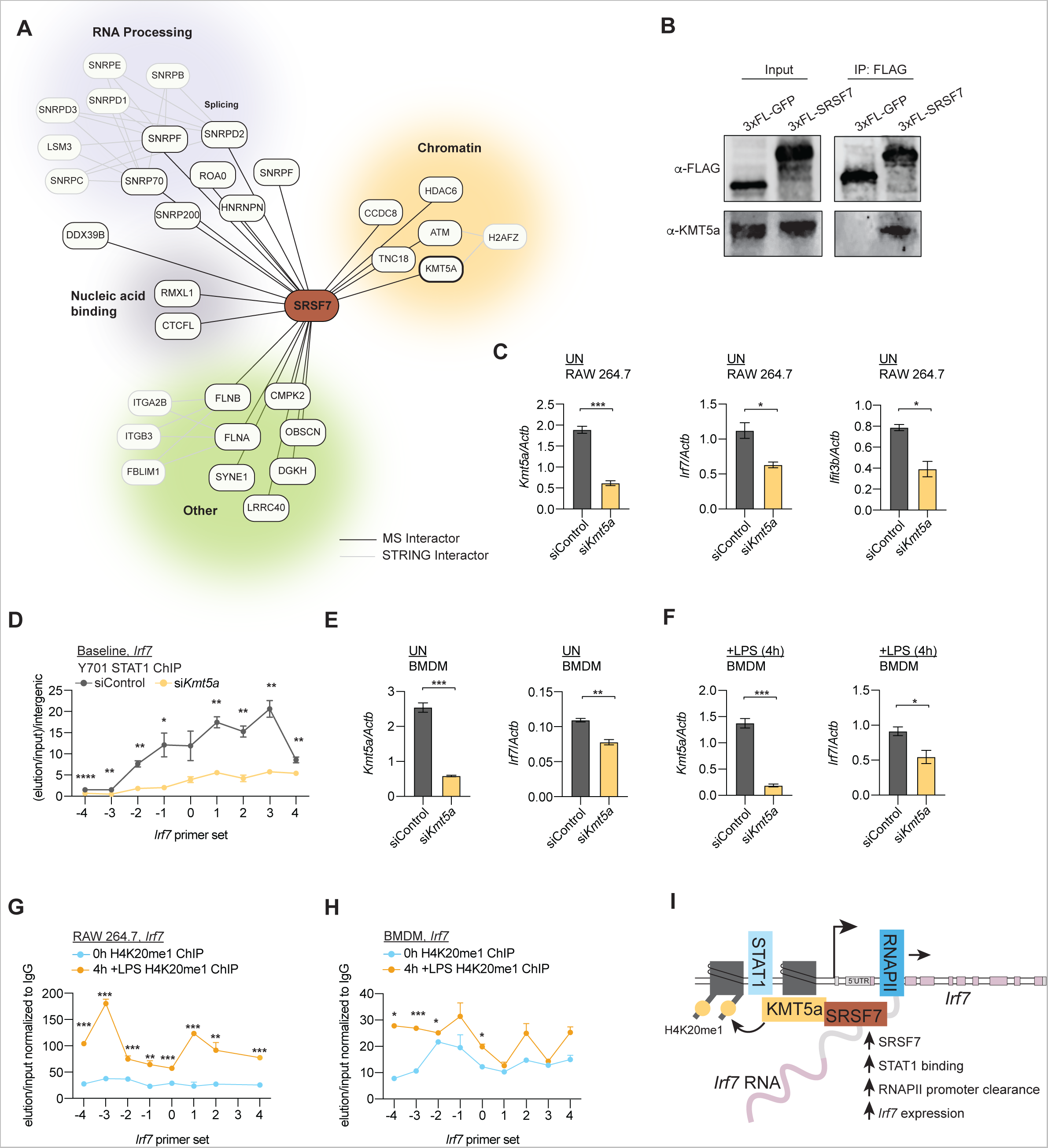
SRSF7 interacts with the H4K20me1 histone methyltransferase KMT5a/SET8 to promote *Irf7* transcription. **A.** Proteins identified as unique interactors of 3xFL-SRSF7 in RAW MΦs at baseline by IP-LC/MS. **B.** Immunoblot of immunoprecipitation of 3xFL-SRSF7 (+DOX 1 µg/ml 22h), 3xFL-GFP (+DOX 1 µg/ml 2h) and endogenous KMT5a/SET8. **C.** RT-qPCR of *Kmt5a, Irf7*, or *Ifit3b* in RAW MΦs transfected with siControl or si*Kmt5a* siRNA at baseline. **D.** Y701 STAT1 ChIP-qPCR at the *Irf7* promoter in siControl or si*Kmt5a/Set8* RAW MΦs at baseline. Data represented as elution/input normalized to that of an intergenic control. **E.** RT-qPCR of *Kmt5a* and *Irf7* in bone marrow derived macrophages (BMDMs) transfected with siControl or si*Kmt5a* siRNA. **F.** As in E but at 4h post-stimulation with LPS (100 ng/ml). **G.** H4K20me1 ChIP-qPCR in RAW MΦs at baseline or 4h post stimulation with 100 ng/ml LPS at the *Irf7* promoter. Data represented as elution/input normalized to IgG. **H.** As in G but in BMDMs. **I.** Proposed model for cooperation between SRSF7 and KMT5a promoting transcriptional activation of *Irf7*. Unless otherwise noted, data are expressed as a mean of three or more biological replicates with error bars depicting SEM. ChIP-qPCRs are shown as the mean of three or more technical replicates with error bars depicting SEM. Statistical significance was determined using two tailed unpaired student’s t test. *=p<0.05, **=p<0.01, ***=p<0.001.

**Figure S6:**
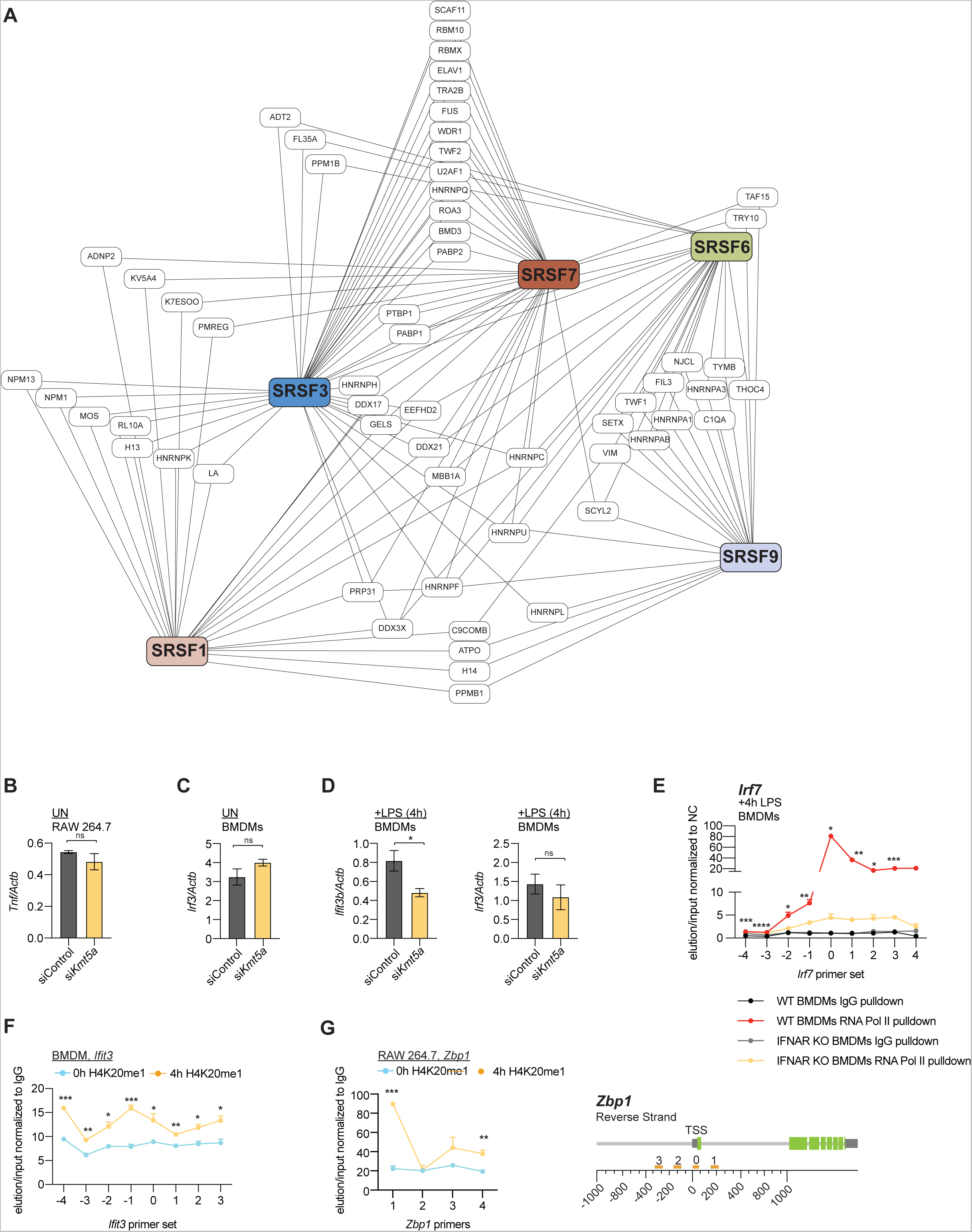
Supplemental data for Figure 6. **A.** Proteins identified as shared interactors of 3xFL-SRSF7, 3xFL-SRSF1, 3xFL-SRSF3, 3xFL-SRSF6, and 3xFL-SRSF9 in RAW MΦs at baseline by IP-LC/MS. **B.** RT-qPCR of *Tnf* in RAW MΦs transfected with siControl or si*Kmt5a* siRNA at baseline. **C.** RT-qPCR of *Irf3* in BMDMs transfected with siControl or si*Kmt5a* siRNA at baseline. **D.** As in C but 4h post-stimulation with 100 ng/ml LPS. **E.** RNA polymerase II ChIP-qPCR 4h post-stimulation with LPS in WT and *Ifnar*-/- BMDMs at the *Irf7* promoter **F.** H4K20me1 ChIP-qPCR at the *Ifit3* promoter in RAW MΦs at baseline or 4h post-stimulation with 100 ng/ml LPS. **G.** As in F at the *Zbp1* promoter. Right: Diagram of the *Zbp1* gene with tiling primers designed around the transcriptional start site (orange). Unless otherwise noted, data are expressed as a mean of three or more biological replicates with error bars depicting SEM. ChIP-qPCRs are shown as the mean of three or more technical replicates with error bars depicting SEM. Statistical significance was determined using two tailed unpaired student’s t test. *=p<0.05, **=p<0.01, ***=p<0.001.

### SRSF7 interacts with the H4K20me1 histone methyltransferase KMT5a/SET8 to promote *Irf7* transcription

ChIP-seq did not reveal a strong consensus sequence for SRSF7 and even weak consensus sequence enrichment was only detected at subsets of genes (**Fig. S5E**). As this argues against recruitment of SRSF7 to the *Irf7* promoter via a specific DNA-encoded motif, we hypothesized that SRSF7 could be brought to/retained at the *Irf7* promoter via one or more protein-protein interactions. To begin to explore this possibility, we generated RAW MΦ cell lines stably expressing 3xFLAG-SRSF1, 3xFLAG-SRSF3, 3xFLAG-SRSF6, 3xFLAG-SRSF9, in addition to the previously described 3xFLAG-SRSF7 and 3xFLAG-GFP cell lines. We performed a series of immunoprecipitation experiments followed by mass spectrometry to identify peptides enriched in each pulldown. Although many protein binding partners were shared between SRSF7 and other SR protein family members (e.g. SR proteins, hnRNPs, RRM domain-containing proteins) (**Fig. S6A**), peptides derived from components of the KMT5a/SET8 histone methyltransferase complex were unique to SRSF7 (**Fig. 6A**). KMT5a is the sole histone methyltransferase responsible for depositing monomethylation at H4K20. H4K20me1 levels are linked to the cell cycle and during mitosis H4K20me1 mediates chromatin condensation (van Nuland and Gozani, 2016). Several studies report enrichment of H4K20me1 at actively transcribed genes (Barski et al., 2007), (Veloso et al., 2014) and it has been implicated in facilitating RNAPII escape from promoter-proximal regions (Nikolaou et al., 2017). We confirmed interaction between 3xFLAG-SRSF7 and KMT5a in macrophages by immunoprecipitation/immunoblot, using antibodies against endogenous KMT5a (**Fig. 6B**).

To determine a role for KMT5a in mediating *Irf7* expression, we silenced its expression via siRNA transfection in RAW MΦs, alongside a non-targeting control. We found that loss of KMT5a dampened *Irf7* transcript levels and those of downstream ISGs, phenocopying SRSF7 knockdown (**Fig. 6C**). Expression of inflammatory genes outside the *Irf7* regulon, like *Tnf*, was not impacted by loss of KMT5a (**Fig. S6B**), hinting at a privileged role for KMT5a in modulating ISGs. Notably, STAT1 failed to associate with the *Irf7* promoter in *Kmt5a* knockdown RAW MΦs (**Fig. 6D**), strongly supporting a role for KMT5a in promoting a pro-transcription chromatin environment at *Irf7*. We also detected lower *Irf7* transcript levels both at rest (**Fig. 6E**) and at 4h post-LPS treatment (**Fig. 6F**) when we knocked down KMT5a in BMDMs. Together, these data implicate KMT5a in stimulating *Irf7* expression in macrophages.

Lastly, we sought to implicate the histone mark deposited by KMT5a, H4K20me1, in activating *Irf7* expression. We found that LPS treatment triggered significant accumulation of H4K20me1 at the *Irf7* promoter in both RAW MΦs (**Fig. 6G**) and BMDMs (**Fig. 6H**) that was concomitant with IFNAR-dependent transcriptional activation and recruitment of RNAPII (**Fig. S6E**). LPS treatment also elevated H4K20me1 levels at ISGs like *Ifit3* and *Zbp1* (**Fig. S6F-G**). Because we did not see SRSF7 enrichment at the promoters of these or other ISGs by ChIP-qPCR or ChIP-seq (**Fig. 5L**), our data argue that H4K20me1 does not recruit SRSF7 to promoters. Instead, our data support a model wherein interaction between SRSF7 and KMT5a confers tight control over the chromatin landscape at *Irf7* promoter, which is required to maximize STAT1 recruitment and promote RNAPII elongation (**Fig. 6I**).

## DISCUSSION

Because of their potential to activate large gene regulons, cells need to exert tight control over transcription factors themselves. This is particularly true for transcription factors that activate inflammatory immune gene expression, which are subject to multiple layers of post-translational regulation (cytoplasmic sequestration, phosphorylation, proteasomal degradation) (Caamano and Hunter, 2002; Jefferies, 2019; Ning et al., 2011; Smale, 2012). Here, we describe a novel role for the splicing regulatory protein SRSF7 in stimulating the type I IFN response by activating transcription of the *Irf7* gene. We demonstrate that enrichment of SRSF7 at the *Irf7* promoter facilitates association of the transcription factor STAT1 and promotes RNAPII elongation and initiate expression of *Irf7*. SRSF7 accomplishes this by establishing a pro-transcription chromatin landscape at the *Irf7* promoter via interaction with the H4K20me1 methyltransferase KMT5a. By controlling IRF7 abundance, SRSF7 plays a privileged role in regulating both antiviral (**Fig. 2Q-T**) and antibacterial (**Fig. 5H**) innate immune responses in macrophages. In addition to implicating SRSF7 in activating *Irf7* expression, this work highlights a critical and underappreciated role for both SRSF7 and IRF7 in maintaining tonic and early interferon signaling in MΦs.

A major finding of this study relates to the ability of SRSF7—a canonical RNA binding protein—to promote transcription of *Irf7*. Over the past two decades, the idea that transcription and RNA processing occur in a linear, stepwise fashion has been repeatedly challenged. It is now generally accepted that many steps in RNA processing (i.e. capping, pre-mRNA splicing, RNA editing) occur co-transcriptionally while nascent transcripts are still bound to RNAPII. Accordingly, splicing can influence transcription and transcription can impact splicing (Tellier et al., 2020). Several studies have linked SR proteins to RNAPII dynamics and co-transcriptional splicing. SRSF2, for example, releases the cyclin-dependent kinase pTEFb from an inhibitory 7SK-containing complex to relieve stalled polymerase at promoters (Ji et al., 2013) and SRSF3 regulates exon inclusion via the C-terminal domain of RNAPII (de la Mata and Kornblihtt, 2006). Although our data suggest an analogous role for SRSF7 in mediating transcription factor binding and RNAPII elongation at the *Irf7* promoter, several aspects of this mechanism still need to be explored.

One outstanding question relates to why we see SRSF7 at certain promoters, like *Irf7*, but not others (**Fig. 6**). Owing to its ability to interact with histone modifying enzymes like KMT5a, it is possible that the local chromatin environment dictates the specificity of SRSF7 promoter enrichment. Our data demonstrate that H4K20me1 levels are significantly enhanced at the promoters of innate genes including *Irf7* following LPS treatment, linking this mark to macrophage activation and innate immune responses for the first time. However, because H4K20me1 is enriched at innate genes like *Ifit3* and *Zbp1* where SRSF7 promoter association was not detected (**Fig. S6F-G**), we currently do not favor a “chromatin recruits SRSF7” model. Instead, we propose that specificity is dictated by sequences encoded in exons/introns of genes downstream of SRSF7-sensitive promoters, such that nascent RNAs serve as the platform for SRSF7 recruitment. The reported consensus sequence for SRSF7 is GA(C/T)GA(C/T) (Cavaloc et al., 1999) and SRSF7 was shown to bind TGA(C/T)C sequences in its own pre-mRNA to regulate poison cassette exon inclusion (Konigs et al., 2020). We can detect several potential SRSF7 bindings sites in the first coding exon of *Irf7* (TGACC). Thus, it is possible that SRSF7’s association at the *Irf7* promoter occurs by virtue of co-transcriptional binding of exonic splicing enhancers (ESEs) in nascent transcripts.

It is also conceivable that SRSF7 associates with the *Irf7* promoter in an RNA-independent, DNA-dependent fashion. SRSF7 is the only SR protein that encodes a Zn-knuckle, and to date, this domain has been shown to enhance the RNA binding activity of SRSF7 (Cavaloc et al., 1999; Cavaloc et al., 1994; Wang et al., 2021)). However, in other proteins, for example TAF1, the largest subunit of the general transcription factor TFIIID, a Zn knuckle encourages DNA binding and promoter occupancy (Curran et al., 2018). It is tempting to speculate that the Zn finger domain in SRSF7 could promote its association directly with DNA. Future experiments designed to implicate the Zn knuckle in SRSF7 association with the *Irf7* promoter and/or test the ability of SRSF7 to directly bind DNA will help resolve the molecular mechanisms of SRSF7 promoter enrichment.

In addition to linking SRSF7 to *Irf7* expression, our studies revealed an unexpected role for SRSF7 in controlling tonic type I IFN expression in resting RAW cells (**Fig. 2A, E**) and MEFs (**Fig. 2M-P**, time 0). These observations challenge the paradigm that IRF7 needs to be dramatically upregulated before it can contribute to type I IFN responses. Indeed, differences in ISG expression were more robust in IRF7 KO RAW MΦs at rest than after LPS stimulation (**Fig. S4B**). These findings suggest that IRF7 is in fact the major mediator of the type I IFN response in resting macrophages, but *both* IRF3 and IRF7 contribute once they are activated downstream of pattern recognition receptor engagement. To date, the role for IRF7 in maintaining tonic type I IFN signaling has not been fully explored. Our RNA-seq data identify a specific regulon of ISGs that is particularly sensitive to loss of IRF7 at baseline (**Fig. S3F, G**). Among the most downregulated genes in resting *Irf7* KO macrophages are several encoding guanylate binding proteins (GBPs) and members of the oligoadenylate synthetase (OAS) family. Regulated expression of these factors at baseline may reflect their role in directly mediating anti-viral and anti-bacterial killing, with low level expression of these ISGs in resting cells providing an early advantage upon infection. By demonstrating a role for IRF7 in maintaining tonic levels of these and other ISGs, our results hint at additional roles for IRF7 in cellular homeostasis and disease that have not been appreciated to date.

Consistent with this idea, growing evidence suggests that IRF7 expression in immune cells is subject to multiple levels of control mediated by RBPs. Both mouse and human IRF7 genes encode very small introns (<100nt), an unusual adaptation in mammalian genomes. Regulated *IRF7* intron retention has been reported in THP-1 cells, wherein intron 2 is efficiently removed upon monocyte-to-macrophage differentiation but not in resting monocytes (Green et al., 2020). In response to treatment with TNF-α and IFN-α, as well as polyI:C transfection, regulated removal of IRF7 intron 4 has been reported and was shown to be controlled by the RBP BUD13, which is enriched on *Irf7* transcripts in BMDMs (Frankiw et al., 2019). Such regulation is proposed to provide innate cells a way to limit the inappropriate expression of genes with high inflammatory potential like IRF7, while leaving cells poised to upregulate these genes when certain signals are received. We did not observe differences in the number of intron reads in *Srsf7* KD vs. SCR RAW cells (**Fig. 3H**), nor did the MAJIQ algorithm identify SRSF7-dependent alternative splicing changes in *Irf7* (**Fig. 1**). We propose that intron retention/alternative splicing of *Irf7* by factors like BUD13 adds an additional layer of regulation on top of the transcriptional control conferred by SRSF7. The fact that IRF7 is subject to regulation at multiple nodes argues that it plays a larger than appreciated role in mediating inflammatory responses in immune cells. Together, these previously published studies and our work presented here, argue that IRF7 is required for basal maintenance and ramping up of type I interferon responses and highlights a preeminent role for RNA binding proteins in regulating innate immune outcomes.

## ACKNOWLEDGEMENTS

We would like the thank members of Patrick and Watson labs for critical reading of the manuscript and invaluable feedback. We would also like to thank Phillip West and members of his lab at Texas A&M School of Medicine for sharing reagents and protocols related to VSV infection. Thanks to Susan Carpenter and her lab for sharing reagents and protocols related to iBMDMs. and to the Genomic and RNA Profiling Core (GARP) at Baylor College of Medicine for RNA-seq and ChIP-seq library generation and sequencing. Funding was provided by NIH/NIGMS to KP (R35GM133720), NIH/NIAID to ROW and KP (R01AI155621), and NIH/NIGMS to HMS (F31GM143893).

## Materials and Methods

### Cell culture

RAW 264.7 macrophage-like cells (RAW MΦs) (TIB-71™ ATCC) (originally isolated from male BALB/c mice), L929 ISRE (Hoffpauir et al., 2020), *Irf7* KO RAW MΦs (this paper), and Lenti-X cells (TaKaRa Bio) were cultured at 37 °C with a humidified atmosphere of 5% CO_2_ in complete media containing high glucose, DMEM (ThermoFisher) with 10% FBS (Millipore) 0.2% HEPES (Thermo Fisher).

### shRNA knockdowns

For RAW MΦs stably expressing scramble knockdown and *Srsf7* knockdown, Lenti-X cells were transfected with a pSICO scramble non-targeting shRNA construct and pSICO *Srsf7* shRNA constructs targeted at exon 3 and exon 4 of *Srsf7* using Polyjet (SignaGen Laboratories). Virus was collected 24 and 48 h post transfection. RAW MΦs were transduced using Lipofectamine 2000 (Thermo Fischer). After 48 h, media was supplemented with Hygromycin (Invitrogen) to select for cells containing the shRNA plasmid.

### *S. enterica* (ser. Typhimurium)

*Salmonella enterica* serovar Typhimurium (SL1344) was obtained from Dr. Denise Monack, Stanford. *S.* Typhimurium were streaked out on LB agar plates and incubated at 37°C overnight.

### *S. enterica* (ser. Typhimurium) infections

RAW MΦs were seeded in 12-well tissue culture-treated plates at a density of 7×10^5^ cells per cell, and cultures of *S.* Typhimurium were grown at 37°C in LB broth containing 0.3M NaCl 16h prior to infection. Bacterial cultures were diluted 1:20 3h prior to infection to reach mid-log phase OD_600_ 0.6-0.8. Mid-log phase bacteria were washed 3x with PBS and pelleted at 5,000 rpm for 3 min. Bacteria were then added to DMEM (Hyclone) with 10% FBS (Sigma) and 2% HEPES (HyClone) at an MOI of 5 for transcripts and MOI of 1 for replication. Infected cells were spun at 1,000 rpm for 5 min then incubated for 10 min at 37 °C prior to gentamycin washes adding fresh media. At indicated times post infection, cells were harvested with TRIzol for RNA isolation described below.

### M. tuberculosis

The Erdman strain was used for all *M. tuberculosis* (Mtb) infections as well as Mtb luxBCADE a luciferase expressing strain. Low passage lab stocks were thawed for each experiment to ensure virulence was preserved. Mtb was cultured in roller bottles at 37 °C in Middlebrook 7H9 broth (BD Biosciences) supplemented with 10% OADC (BD Biosciences), 0.5% glycerol (Fisher), and 0.1% Tween-80 (Fisher). All work with Mtb was performed under Biosafety Level 3 containment using procedures approved by the Texas A&M University Institutional Biosafety Committee.

### Mtb in vitro infections

To prepare the inoculum, bacteria were grown to mid log phase (OD 0.6–0.8), spun at low speed (500 rcf) to remove clumps, and then pelleted and washed with PBS 2X. Resuspended bacteria were sonicated and spun at low speed again to further remove clumps. Then Mtb was then diluted in DMEM plus 10% horse serum (Gibco) and added to cells at a multiplicity of infection (MOI) of 10 for RNA and protein analysis, a MOI of 5 for cell death studies, and a MOI of 1 for bacterial growth assays. The day before the infection, RAW MΦs were plated on 12-well tissue culture–treated plates at a density of 3×10^5^ cells per well or plated in corning 96 well black plates at 2.5×10^4^ cells/well and allowed to rest overnight. Cells were spun with bacteria for 10 min at 1,000 rcf to synchronize infection, washed 2X with PBS, and then incubated in fresh media. Where applicable, RNA was harvested from infected cells using 0.25 ml TRIzol reagent at each time point. For bacterial growth assays, RAW MΦs were plated on 12-well tissue culture– treated plates at a density of 2.5×10^5^ cells per well. Luminescence was read for *M. tuberculosis luxBCADE* by lysing in 250 μl 0.5% Triton X-100 and dividing sample into duplicate wells of a 96-well white bottomed plate (Costar). Luminescence was measured and normalized to background using the luminescence feature of the INFINITE 200 PRO (Tecan) at 0, 48, 72 and 92 h post infection.

### Vesicular stomatitis virus

Recombinant *Vesicular stomatitis* virus (VSV; Indiana serotype) containing a GFP reporter cloned downstream of the VSV G-glycoprotein (VSV-G/GFP) was obtained collaboratively from Dr. John Rose at Yale School of Medicine though Dr. A. Phillip West at Texas A&M Health Science Center.

### Vesicular stomatitis Virus (VSV) infections

RAW MΦs were seeded in 12-well tissue culture-treated plates at a density of 7×10^5^ cells per well 16h before infection. The next day cells were infected with VSV-GFP virus (Dalton and Rose, 2001) at an MOI of 0.1 in serum-free DMEM (HyClone). After 1h of incubation with media containing virus, supernatant was removed, and fresh DMEM plus 10% FBS (Millipore) was added to each well. At indicated times post infection, cells were harvested with TRIzol for RNA isolation described below.

### M. musculus

Mice used in this study were C57BL6/J (Stock #00064) initially purchased from Jackson Labratory and afterward maintained with filial breeding. All mice used in experiments were compared to age- and sex-matched controls. Littermates were used for experiments. Mice used to generate BMDMs were males between 10-16 weeks old. Embryos used to make primary MEFs were 14.5 days post-coitum. All animals were housed, bred, and studied at Texas A&M Health Science Center under approved Institutional Care and Use Committee guidelines. All experiments for this study were reviewed and approved by the Texas A&M University Institutional Animal Care and Use Committee (AUP# 2019–0083). Mice were fed 4% standard chow and were kept on a 12 h light/dark cycle and provided food and water *ad libitum*. Mice were group housed (maximum 5 per cage) by sex on ventilated racks in temperature-controlled rooms.

### Primary cell culture

Bone marrow derived macrophages (BMDMs) were differentiated from bone marrow (BM) cells isolated by washing mouse femurs with 10 ml DMEM 1 mM sodium pyruvate. Cells were then centrifuged for 5 min at 400 rcf and resuspended in BMDM media (DMEM, 20% FBS (Millipore), 1 mM sodium pyruvate (Lonza), 10% MCSF conditioned media (Watson lab)). BM cells were counted and plated at 5×10^6^ cells per 15 cm non-tissue culture treated dishes in 30 ml complete BMDM media. Cells were fed with an additional 15 ml of BMDM media on day 3. Cells were harvested on day 7 with 1X PBS EDTA (Lonza).

Mouse embryonic fibroblasts (MEFs) were isolated from embryos. Briefly, embryos were dissected from yolk sacs, washed two times with cold 1X PBS, decapitated, and stripped of peritoneal contents. Headless embryos were disaggregated in cold 0.05% trypsin-EDTA (Lonza) and incubated on ice for 20 min., followed by incubation at 37 °C for an additional 20 min. Cells were then DNase treated with 4 ml disaggregation media (DMEM, 10% FBS, 100 μg/ml DNASE I (Worthington)) for 20 min at 37 °C. Isolated supernatants were spun down at 1000 rpm for 5 min. Cells were resuspended in complete MEF media (DMEM, 10% FBS, 1 mM sodium pyruvate), and plated in 15 cm tissue culture-treated dishes 1 dish per embryo in 25 ml of media. MEFs were allowed to expand for 2-3 days before harvest with 0.05% trypsin-EDTA (Lonza). iBMDMs were immortalized with J2 virus to generate iBMDMs.

### siRNA knockdowns in MEFs

Knockdown of mRNA transcripts was performed by plating 3×10^6^ MEFs in 10 cm plates and rested overnight. The following day complete media was replaced with 5 ml fresh complete media 30 min prior to transfection. Cells were transfected using Fugene SI reagent 10 μM of siRNA stock against either *Srsf7* (ThermoFisher, s105011) or negative control, Silencer® Select Negative Control #1 (4390843). Cells were incubated for 48h in transfection media at 37 °C with 5% CO_2_ prior to downstream experiments.

### siRNA knockdowns in BMDMs

The day prior to transfection, 6×10^5^ BMDMs were seeded into 12-well plates. The day of transfection, 30nM of siRNA against *Srsf7* (ThermoFisher, s105011), against *Kmt5a/Setd8* (ThermoFisher, s85986) or negative control, Silencer® Select Negative Control #1 (4390843) were mixed with 3 µl of LipoRNAiMAX (13778150, Thermo Fisher Scientific), incubated for 10m at room temperature, and then added into wells with 500 µl plain DMEM. Transfection proceeded for 6h and then wells were replenished with complete media.

### Tetracycline inducible cell line generation

For generation of tetracycline inducible RAW MΦs, pLenti CMV rtTA3 Blast (Addgene w756-1) stably expressing clonal RAW MΦs were transduced with pLenti CMV Puro DEST (Addgene w118-1) constructs containing 3xFL-GFP, 3xFL-SRSF7, and al 3xFL-SRSF7 phosphorylation mutants. After 48h, construct containing cells were selected through addition of puromycin (Invivogen). 1 μg/ml doxycycline (Sigma-Aldrich) treatment was used to activate construct expression.

### Cell stimulation for immune activation

7.5×10^5^ RAW MΦs were plated on 12-well tissue-culture treated plates 16h before cellular stimulation assays. The next day, cells were treated with lipopolysaccharide (LPS) from *E. coli* (Invivogen) at 100 ng/ml, or ISD (IDT, annealed in house) at 1 μg/ml, or IFN-β (PBL Assay Science) at 200 I/U per ml for the respective time points. Cells were collected for RNA isolation using Trizol reagent.

### *Srsf7* KD mRNA sequencing and analysis

As reported in Wagner, Scott and West et al. 2021, RNA-seq analysis was performed on RAW MΦs containing shRNA knockdowns of *Srsf6*, *Srsf2*, *Srsf1*, *Srsf7*, and *Srsf9* compared to SCR control with biological triplicates of each cell line. RNA-Seq and library preparation was performed by Texas A&M AgriLife Genomics and Bioinformatics Service. RNA samples were sequenced on Illumina 4000 using 2 x 150-bp paired-end reads. Raw reads were filtered and trimmed and Fastq data was mapped to the Mus musculus Reference genome (RefSeq) using CLC Genomics Workbench 8.0.1. Differential expression analyses were performed using CLC Genomics Workbench. Relative transcript expression was calculated by counting Reads Per Kilobase of exon model per Million mapped reads (RPKM). Statistical significance was determined by the EDGE test via CLC Genomics Workbench. Differentially expressed genes (DEGs) were selected as those with p value threshold < 0.05. Critical pathways affected by SR protein knockdown compared to control RAW MΦs, were analyzed by canonical pathway analysis using Ingenuity Pathway Analysis software from QIAGEN Bioinformatics. Genes that were differentially expressed with a p-value < 0.05 from our RNA-SEQ analysis were used as input. Integrated genomics viewer (IGV) visualization tracks were generated using the RNA-SEQ filtered and trimmed reads.

### *Irf7* KO mRNA sequencing and analysis

RNA-seq analysis was performed on control GFP KO or *Irf7* KO1 RAW MΦs in biological triplicate of each cell line. Data was analyzed by ROSALIND® (https://rosalind.bio/), with a HyperScale architecture developed by ROSALIND, Inc. (San Diego, CA). Reads were trimmed using cutadapt. Quality scores were assessed using FastQC. Reads were aligned to the Mus musculus genome build GRCm39 using STAR^3^. Individual sample reads were quantified using HTseq^4^ and normalized via Relative Log Expression (RLE) using DESeq2 R library. Read Distribution percentages, violin plots, identity heatmaps, and sample MDS plots were generated as part of the QC step using RSeQC. DEseq2 was also used to calculate fold changes and p-values and perform optional covariate correction. Clustering of genes for the final heatmap of differentially expressed genes was done using the PAM (Partitioning Around Medoids) method using the fpc R library. Hypergeometric distribution was used to analyze the enrichment of pathways, gene ontology, domain structure, and other ontologies. The topGO R library was used to determine local similarities and dependencies between GO terms in order to perform Elim pruning correction.

### Alternative splicing analysis

Modeling Alternative Junction Inclusion Quantification (MAJIQ) and the Voila visualization package were used to model alternative splicing events with default parameters described by Vaquero-Garcia et al., 2016. Junction-spanning reads were used to construct splice graphs for transcripts by using the RefSeq annotation supplemented with de-novo (junctions not in the RefSeq transcriptome database but had sufficient evidence in the RNAseq data) detected junctions. Resulting splice graphs were analyzed for all identified local splice variations (LSVs). MAJIQ then quantified expected percent spliced in (PSI) value in control and knockdown samples for every junction in each LSV. Resulted viewed on VOILA were filtered for highest confidence change in LSVs (one or more junctions had at least a 95% probability of expected dPSI of at least an absolute value of 0.2 PSI units between control and knockdown.

### RNA isolation and qPCR analysis

For transcript analysis, cells and tissue were harvested in TRIzol and RNA was isolated using Direct-zol RNA Miniprep kits (Zymo Research) with 1 h DNase treatment. cDNA was synthesized with iScript cDNA Synthesis Kit (Bio-Rad). CDNA was diluted as appropriate for treatment, and a pool of cDNA was used to make a 1:3 standard curve with each standard sample diluted 1:5 to produce a linear curve. RT-qPCR was performed using Power-Up SYBR Green Master Mix (Thermo Fisher) using a Quant Studio Flex6 (Applied Biosystems). Samples were run in triplicate wells in a 384-well plate. Averages of the raw values were normalized to average values for the same sample with the control gene, *Actb*.

### Semiquantitative qPCR analysis

cDNA was synthesized by iScript cDNA Synthesis Kit (Bio-Rad) using an extended 1h amplification. Q5 high fidelity 2X Master mix (New England Biolabs) was used for PCR amplification using targeted primers. Loading dye was added to PCR products and samples were run on 2% agarose gel containing ethidium bromide at 100 volts for 1 h. Gels were imaged on LiCOR Odyssey Fc Dual-Mode Imaging System and bands were quantified.

### Chromatin immunoprepitation

6×10^6^ RAW MΦs or BMDMs were seeded in a 10 cm dish, and 16h later stimulated for the indicated time. After stimulation cells were scraped, transferred to a 50 ml conical and fixed with 1% (v/v) formaldehyde for 10 min with gentle shaking. Formaldehyde was quenched with 0.125 M glycine for 5 min. Cells were pelleted and washed 1x with PBS, and then cells were lysed in 1 ml of ChIP lysis buffer (150 mM NaCl, 5 mM EDTA, 0.5% NP-40 and 1% Triton X-100, 50 mM Tris-HCl, and protease/phosphatase inhibitor (added immediately before lysis) pH 7.5). Lysates were flash frozen, or ChIP proceeded by sonicating with for 40 min (30 sec on/off 10 min cycles) in a Bioruptor UCD-200 (Diagenode). Lysates were cleared and 10% taken for input samples.

For RNA polymerase II, Y701 STAT1, IgG (isotype control) or H4K20me1 samples: 2 µg of the anti-pol II (Active Motif) or 2 µg of isotype control (IgG, Cell Signaling), 5ul Y701 STAT1 (Cell Signaling), or 2 µg H4K20me1 (Invitrogen) (were incubated overnight at 4 with rotation. The following day, Pierce beads (Invitrogen) were washed 3x in ChIP buffer, blocked for 30 min with 100 µg salmon sperm DNA (Invitrogen) and 0.5 mg BSA, and washed once more with ChIP lysis buffer. Blocked beads were incubated with cleared chromatin immunocomplexes for 1h at 4 °C with rotation. Beads were washed five times with ChIP buffer and then eluted with 250 µl of elution buffer (1% SDS and 0.1 M NaHCO_3_) at 65 °C for 15 mins. The eluate and input were centrifuged and reverse the cross-links by adding 0.4 M NaCl for overnight at 55 °C. The next day, 1 µl of RNASE A (10 mg/ml, ThermoFisher) and 2 µl proteinase K (20 mg/ml, ThermoFisher) were added to the samples and incubated for 1h at 55 °C while shaking. Samples were then subjected to Genomic Clean up and Concentrator (Zymo) to extract DNA. Purified input and IP DNA were analyzed by qRT-PCR using primers specific for the promoter region of target genes. Results were normalized to input. Data is presented as fold enrichment relative to intergenic control. <u>Flag ChIP</u>: As above except cleared lysates were added to blocked (as above) M2 anti-3xFLAG beads and protocol proceeded as above.

### 3x-FLAG ChIP-seq

Data was analyzed by ROSALIND® (https://rosalind.bio/), with a HyperScale architecture developed by ROSALIND, Inc. (San Diego, CA). Reads were trimmed using cutadapt^1^. Quality scores were assessed using FastQC^2^. Reads were aligned to the Mus musculus genome build mm10 using bowtie2^3^. Per-sample quality assessment plots were generated with HOMER^4^ and Mosaics^5^. Peaks were called using MACS2^6^ (with input/IgG controls background subtracted, if provided). Peak overlaps and differential binding were calculated using the DiffBind R library^7^. Differential binding was calculated at gene promoter sites. Read distribution percentages, identity heatmaps, and FRiP plots were generated as part of the QC step using ChIPQC R library^8^ and HOMER. HOMER was also used to generate known and de novo motifs and perform functional enrichment analysis of pathways, gene ontology, domain structure and other ontologies.

### CRISPR/Cas9 KO of *Irf7*

Guide RNAs were generated to target the 3^rd^ exon of *Irf7* using the Broad online tool (https://portals.broadinstitute.org/gppx/crispick/public) and synthesized by IDT. Primers were cloned into digested vector by phosphorylating, annealing, and ligating primers into LentiGuide-Puro (Addgene, plasmid 52963). Colonies were validated by Sanger sequencing using pLK0.1/hU6 promoter primer (Eton Biosciences, San Diego, CA). Lentivirus with sgRNAs were produced and used to transduce RAW MΦs stably expressing FL-Cas9 (Addgene, plasmid 52962). After selection with 5 µg/ml puromycin knockout efficiency was analyzed by immunoblot and then sequencing for mutants by amplifying a 300 bp region of genomic DNA around exon 3. These PCR fragments were sequenced (as above) and compared to controls using TIDE analysis. The two best, most efficient sgRNA populations were clonally selected, expanded, and assayed for clones with total KO of IRF7 by immunoblot.

### Protein quantification by immunoblot

Cells were washed with PBS and lysed in 1X RIPA buffer with protease and phosphatase EDTA free inhibitors (Thermo Scientific), with the addition of 1 U/ml Benzonase (Millipore) to degrade genomic DNA. Proteins were separated by SDS-PAGE on Any kD mini-PROTEAN TGX precast gel (Bio-Rad) and transferred to 0.45μm nitrocellulose membranes (GE Healthcare). Membranes were blocked for 1h at RT in TBS +5% BSA. Blots were incubated overnight at 4 °C with the following primary antibodies: β-ACTIN (Abcam, 1:5000), Y701 STAT1 (Cell Signaling, 1:1000), STAT1 (Cell Signaling, 1:1000), IRF7 (Cell Signaling, 1:1000), SRSF7 (Bethyl 1:3000) ZBP1 (Santa Cruz Biotechnologies, 1:1000), p-IRF3(S396) (Cell Signaling, 1:1000); IRF3 (Bethyl, 1:1000); ANTI-FLAG M2 (Sigma Aldrich, 1:5000); Membranes were washed 3 x 5 min in PBS-Tween20 and incubated with appropriate secondary antibodies (LI-COR) for 1h at RT prior to imaging on a LiCOR Odyssey Fc Dual-Mode Imaging System.

### Quantitation and Statistical Analysis

Statistical analysis of data was performed using GraphPad Prism software. Two-tailed unpaired Student’s t tests were used for statistical analyses, and unless otherwise noted, all results are representative of at least three biological samples (mean +/- SEM (n = 3 per group)).

